# A novel miR-99b-5p-*Zbp1* pathway in microglia contributes to the pathogenesis of schizophrenia

**DOI:** 10.1101/2023.03.21.533602

**Authors:** Lalit Kaurani, Md Rezaul Islam, Urs Heilbronner, Dennis M. Krüger, Jiayin Zhou, Aditi Methi, Judith Strauss, Ranjit Pradhan, Susanne Burkhardt, Tonatiuh Pena, Lena Erlebach, Anika Bühler, Monika Budde, Fanny Senner, Mojtaba Oraki Kohshour, Eva C. Schulte, Max Schmauß, Eva Z. Reininghaus, Georg Juckel, Deborah Kronenberg-Versteeg, Ivana Delalle, Francesca Odoardi, Alexander Flügel, Thomas G. Schulze, Peter Falkai, Farahnaz Sananbenesi, Andre Fischer

## Abstract

Schizophrenia is a psychiatric disorder that is still not readily treatable. Pharmaceutical advances in the treatment of schizophrenia have mainly focused on the protein coding part of the human genome. However, the vast majority of the human transcriptome consists of non-coding RNAs. MicroRNAs are small non-coding RNAs that control the transcriptome at the systems level. In the present study we analyzed the microRNAome in blood and postmortem brains of controls and schizophrenia patients and found that miR-99b-5p was downregulated in both the prefrontal cortex and blood of patients. At the mechanistic level we show that inhibition of miR-99b-5p leads to schizophrenia-like phenotypes in mice and induced inflammatory processes in microglia linked to synaptic pruning. The miR-99b-5p-mediated inflammatory response in microglia depended on *Z-DNA binding protein 1* (*Zbp1*) which we identified as a novel miR-99b-5p target. Antisense oligos (ASOs) against *Zbp1* ameliorated the pathological phenotypes caused by miR-99b-5p inhibition. In conclusion, we report a novel miR-99b-5p-*Zbp1* pathway in microglia that contributes to the pathogenesis of schizophrenia. Our data suggest that strategies to increase the levels of miR-99b-5p or inhibit *Zbp1* could become a novel therapeutic strategy.

## Introduction

Schizophrenia (SZ) is a devastating psychiatric disorder, and the difficulties involved in treating and managing it make it one of the ten most expensive disorders for health care systems worldwide [1] [2]. SZ is believed to evolve on the background of complex genome-environment interactions that alter the cellular homeostasis as well as the structural plasticity of brain cells. Thus, genetic predisposition and environmental risk factors seem to affect processes that eventually contribute to the manifestation of clinical symptoms [3] [4] [5]. Despite the available pharmacological and non-pharmacological treatment options, a significant number of patients do not benefit from these treatments in the long-term, underscoring the need for novel and potentially stratified therapeutic approaches [6]. So far, drug development has focused on the human transcriptome that encodes proteins, but the success of this approach is limited [7]. However, most of the transcriptome consists of non-coding RNAs (ncRNAs) which are recognized as key regulators of cellular functions [8]. Therefore, RNA therapeutics represent an emerging concept that may expand current therapeutic strategies focused on the protein-coding part of our genome [9] [10]. RNA therapies utilize, for example, antisense oligonucleotides (ASOs), siRNA, microRNA (miR) mimics or corresponding anti-miRs to control the expression of genes and proteins implicated in disease onset and progression [11] [9]. Of particular interest are miRs, which are 19-22 nucleotide-long RNA molecules that regulate protein homeostasis via binding to target mRNAs, leading either to their degradation or reduced translation [12]. miRs have been intensively studied as biomarkers and therapeutic targets in cancer [11] and cardiac diseases [13]. There is also emerging evidence from genetic studies in humans as well as functional data from mouse models that miRs play a role in CNS diseases including SZ [14] [15] [16] [17]. In addition, several studies reported changes in miR expression in blood samples of SZ patients using either qPCR analysis of selected targets or genome-wide approaches. The current findings have been summarized in several review articles [18] [19] [20]. Despite this progress, there are still only few reports on the function of candidate miRs [21]. Nevertheless, analysis of miRs in liquid biopsies is highly valuable because one miR can affect many target genes, and thus changes in miR expression can indicate the presence of multiple pathologies [22] [14] [23]. Moreover, miRs also participate in inter-organ communication [24] [25], suggesting that alterations of miR expression in liquid biopsies may inform about relevant pathological processes in other organs, including the brain. This is important since the analysis of the molecular processes underlying neuropsychiatric diseases in post-mortem human brain tissue is challenging because it might be affected e.g. by peri-mortem events or the timing of post-mortem tissue sampling. Furthermore, the onset of the disease often precedes tissue collection by decades. In contrast, liquid biopsies such as blood samples are easy to collect on the premise that molecular changes in blood mirror changes in the brain. In this context, the analysis of the microRNAome in liquid biopsies could be a suitable approach to identify candidate microRNAs that may play a role in the onset and progression of SZ.

In the present study we performed small RNA sequencing in blood samples of control participants (n = 331) and schizophrenia patients (n = 242) of the PsyCourse Study [26] (http://www.psycourse.de/). By cross-correlating our findings with data from post-mortem human brain tissue, we identified miR-99b-5p as a promising biomarker candidate that is decreased in blood and in the prefrontal cortex of SZ patients, and correlates with disease phenotypes. Furthermore, we found decreased levels of miR-99b-5p in the prefrontal cortex of mice to elicit SZ-like phenotypes and activate pathways linked to innate immunity. In line with these observations, inhibition of miR-99b-5p in microglia increased phagocytosis and reduced the number of synapses. Finally, we were able to demonstrate that this effect is controlled by the miR-99b-5p target gene *Zbp1*, an upstream regulator of innate immunity [27]. Taken together, our data suggest that targeting miR-99b-5p or its target *Zbp1* could provide a novel approach towards the treatment of SZ patients.

## Results

### miR-99b-5p expression is decreased in SZ patients

To identify microRNAs that play a role in the pathogenesis of SZ, we conducted small RNA sequencing of blood samples obtained from 573 participants of the PsyCourse Study (http://www.psycourse.de/). We analyzed 331 healthy controls and 242 SZ patients **(Fig. 1a, Fig. EV1, supplemental table 1)** [26]. After data normalization and correction for confounding factors, we performed a weighted gene co-expression network analysis (WGCNA) and detected 8 co-expression modules that significantly differed between groups. Three were significantly decreased **(Fig. 1b)** while 5 were increased **(Fig. 1c)** in SZ patients **(supplemental table 2)**. The turquoise, yellow, blue and red modules displayed the most significant differences among the groups (*P* < 0.0001). We then investigated whether the expression levels of any of these modules correlated with the clinical phenotypes defined by the positive and negative syndrome rating scale (PANSS) and Beck depression inventory (BDI-II), both of which are decreased in SZ patients, and/or the global assessment of functioning (GAF) score, which is increased in these patients **(Fig. EV1)**. Out of the 8 co-expression modules the turquoise, yellow, blue and red modules were significantly correlated to all disease phenotypes. The turquoise module exhibited a significant negative correlation to the PANSS and BDI-II score and a positive correlation with the GAF score **(Fig. 1d)**, which is in line with the decreased expression of this module in SZ. The green, yellow, blue and red modules were positively correlated to the PANSS and BDI-II scores and exhibited a negative correlation with the GAF score, which is in agreement with their increased expression in SZ **(Fig. 1d)**. The pink module was negatively correlated to the PANSS scores and positively correlated to the GAF, which is also in agreement with its decreased expression in SZ patients. However, the pink module was not correlated to the BDI-II score **(Fig. 1d)**. The black and the brown modules were not significantly correlated any of the analyzed phenotypes. Taken together, these data suggest that especially the microRNAs present in turquoise, pink, green, blue, yellow and red modules warrant further analysis. When we subjected the confirmed targets of the microRNAs present within each of these modules to a GO-term analysis, we observed that most modules were linked to inflammatory processes, which is in agreement with a suggested role of neuro-inflammation in the pathogenesis of SZ [28] **(Fig. EV2; Supplemental table 3)**. While WGCNA is a suitable first approach to identify groups of candidate microRNAs, we also performed a differential expression analysis to directly compare the microRNAome in control vs SZ patients. We found 59 microRNA that were significantly increased in SZ patients while 34 microRNAs were decreased **(Fig 1e, Supplemental table 4).**

**Figure 1:**
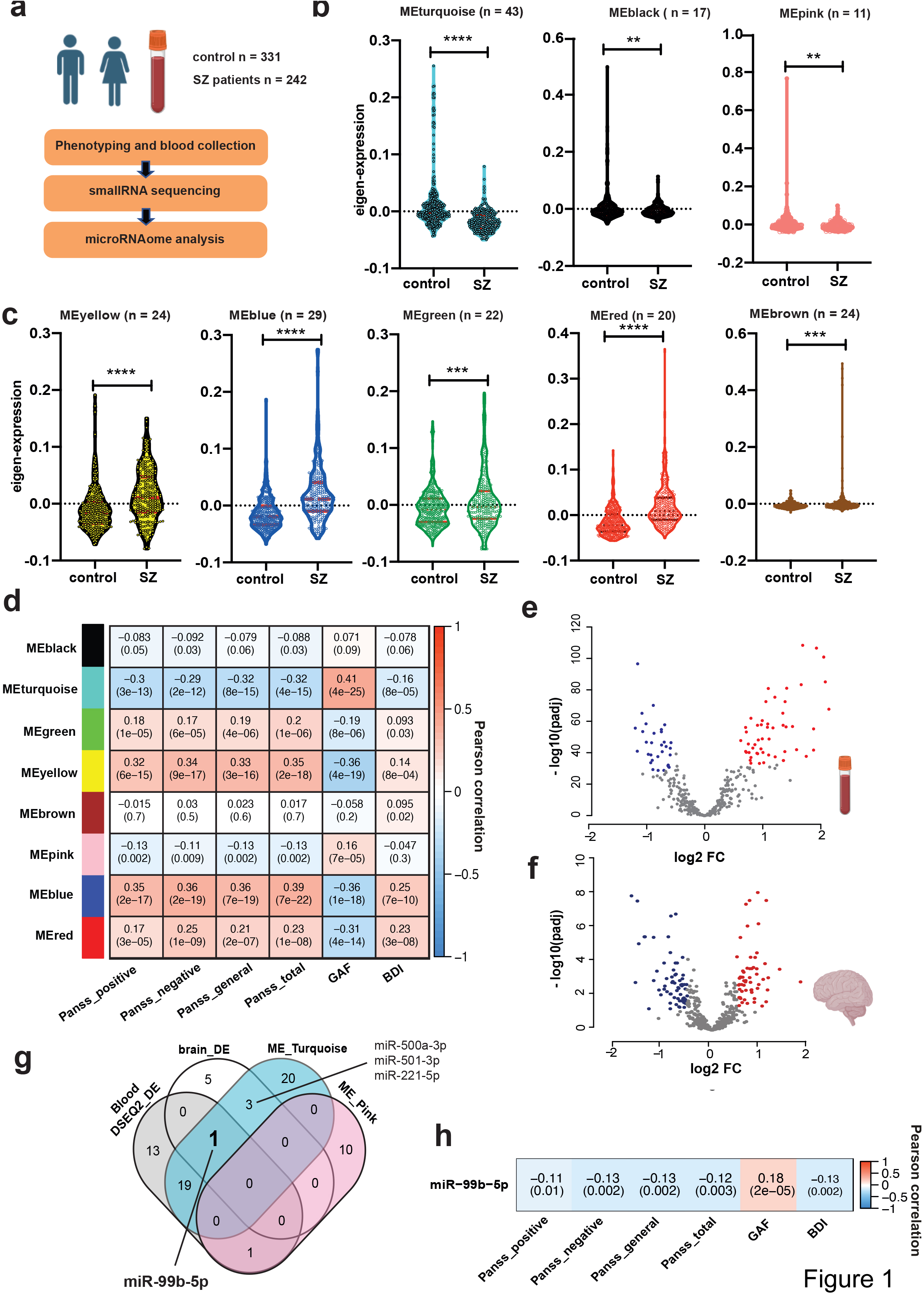
Identification of microRNAs that play a role in the pathogenesis of SZ. **a.** Experimental scheme. **b.** Violin plots showing the results of WGCNA analysis. Depicted is the comparison of the eigenexpression in of the 3 co-expression modules in SZ patients and in control, showing a decrease in SZ patients. **c.** Violin plots showing the results of WGCNA analysis. The eigenexpression of the 5 co-expression modules was higher in SZ patients than in controls (for b & c: unpaired t test; **P < 0.01, ***P < 0.001; ****P < 0.0001, a P value < 0.01 was considered as significant). **d.** Heat map showing the correlation of the eigenexpression of the co-expression modules shown in (a) and (b) with the corresponding clinical phenotypes. The numbers in each rectangle represent the correlation (upper number) and the corresponding p-value (lower number). A P value < 0.01 was considered as significant. **e.** Volcano plot depicting the results of the differential expression analysis when comparing SZ patients and controls shown in (a). **f.** Volcano plot demonstrating the results of the differential expression when comparing postmortem brain samples from SZ patients (n=13) and controls (n=17). **g.** Venn diagram comparing the microRNAs detected in blood samples when performing differential expression analysis (Blood DESEQ2_DE), the microRNAs of the ME_Turquoise and ME_Pink co-expression modules and the microRNAs differentially expressed when comparing postmortem brain tissue (brain_DE). miR-99b-5p is the only microRNA decreased in all comparisons. **h.** Heat map showing the correlation of miR-99b-5p expression levels to the clinical phenotypes for the individuals as analyzed in (a). The numbers in each rectangle represent the correlation (upper number) and the corresponding p-value (lower number).

With the aim to further refine the list of candidate microRNAs that may play a role in the pathogenesis of SZ, we performed small RNA sequencing from the prefrontal cortex of SZ patients (n=13) and controls (n=17). We detected 36 microRNAs that were significantly decreased in SZ patients (FDR < 0,01, log2FC > 1) and 32 that were significantly upregulated **(Fig. 1f, supplemental table 5)**. Next, we examined whether any of these microRNAs are also found among the differentially expressed microRNAs that were altered in blood samples when compared via differential expression analysis, and within the co-expression modules decreased in SZ patients. Three miRs of the yellow cluster were significantly increased in the brain and in the blood when analyzed via differential expression analysis. In addition, one miR of the blue and one of the green clusters were also increased in brain and blood. These were miR-101-3p, miR-378a-3p, miR-21-5p, miR-192-5p and miR-103a-3p **(supplemental table 6)**. MiR-21-3p has been associated with SZ while the other 4 miRs have been studied in the context of other neuropsychiatric or neurodegenerative diseases **(supplemental table 6)**. When analyzing the down-regulated miRs we found four microRNAs, namely miR-500a-3p, miR-501-3p, miR-221-5p and miR-99b-5p, that were detected in the turquoise module and were also decreased in the postmortem brain of SZ patients **(Fig. 1g)**. Of these 4 microRNAs miR-501-3p has been recently linked to schizophrenia [21] **(supplemental table 6)** while miR-99b-5p was the only candidate that was part of a significantly downregulated co-expression module and was significantly downregulated in the brain and blood of SZ patients when analyzed via differential expression analysis.

In summary, our data reveal a number of interesting candidate miRs such as miR-501-3p and miR-21-5p that have been already linked to SZ in previous studies [29] [21]. Most of the other miRs have been detected in the context of other brain diseases including Alzheimer’s disease (AD), Major depression (MD) or Amytrophic lateral sclerosis (ALS) **(supplemental table 6)**. Expect for miR-500a-3p and miR-221-5p, the expression of all other candidate miRs have is significantly correlated to the PANSSs, GAF and BDI-II scores **(Fig. 1h, Fig. EV3)**. While all of these candidate miRs would warrant further functional analysis in the context of SZ, we decided to focus on miR-99b-5p since it has not been linked to any brain disease yet and comparatively little is known about this miR in general.

### Decreasing miR-99b-5p leads to SZ-like phenotypes in mice and the upregulation of genes linked to inflammatory processes

The role of miR-99b-5p in the brain has not been intensively studied and thus no data are available in the context of neuropsychiatric diseases such as SZ, making it a novel candidate in need of further evaluation. Before performing mechanistic studies, we decided to employ mice in a model system to test the hypothesis that decreased expression of miR-99b-5p is causatively linked to the development of SZ-like phenotypes. Therefore, we generated lipid nanoparticles (LNPs) containing locked nucleic acid (LNA) representing miR-99b-5p inhibitors (anti-miR99b) and injected these into the prefrontal cortex (PFC) of mice. LNAs representing a scrambled sequence were used as controls (sc-control). MiR-99b-5p levels were significantly decreased in the PFC when measured 5 or 10 days after the injections **(Fig 2a)**. To test schizophrenia-like behaviors in animals, we injected either anti-miR-99b or sc-control to the PFC of mice. Explorative behavior measured in the open field test was similar in all groups **(Fig 2b)**. However, anti-miR-99b-treated mice spent less time in the center of the open field, which is indicative of increased anxiety **(Fig 2c)**, a phenotype commonly observed in schizophrenia patients [30]. We also analyzed anxiety behavior in the elevated plus maze test. Anti-miR-99b treated mice spent less time in the open arms, which indicates increased anxiety **(Fig 2d)**. Another valid animal model to test schizophrenia-like behavior in rodents is the pre-pulse inhibition of the startle response (PPI) which is used to measure sensory-gating function [31]. PPI is impaired in SZ patients, can easily be assayed in mice and is impaired in mouse models for SZ [32] [33]. We observed that mice injected with anti-miR-99b displayed significantly impaired PPI responses when compared to the sc-control group **(Fig 2e)**. The basic startle response was unaffected **(Fig. 2f)**, suggesting that decreasing the levels of miR-99b-5p in the PFC of mice can lead to SZ-like phenotypes.

**Figure 2:**
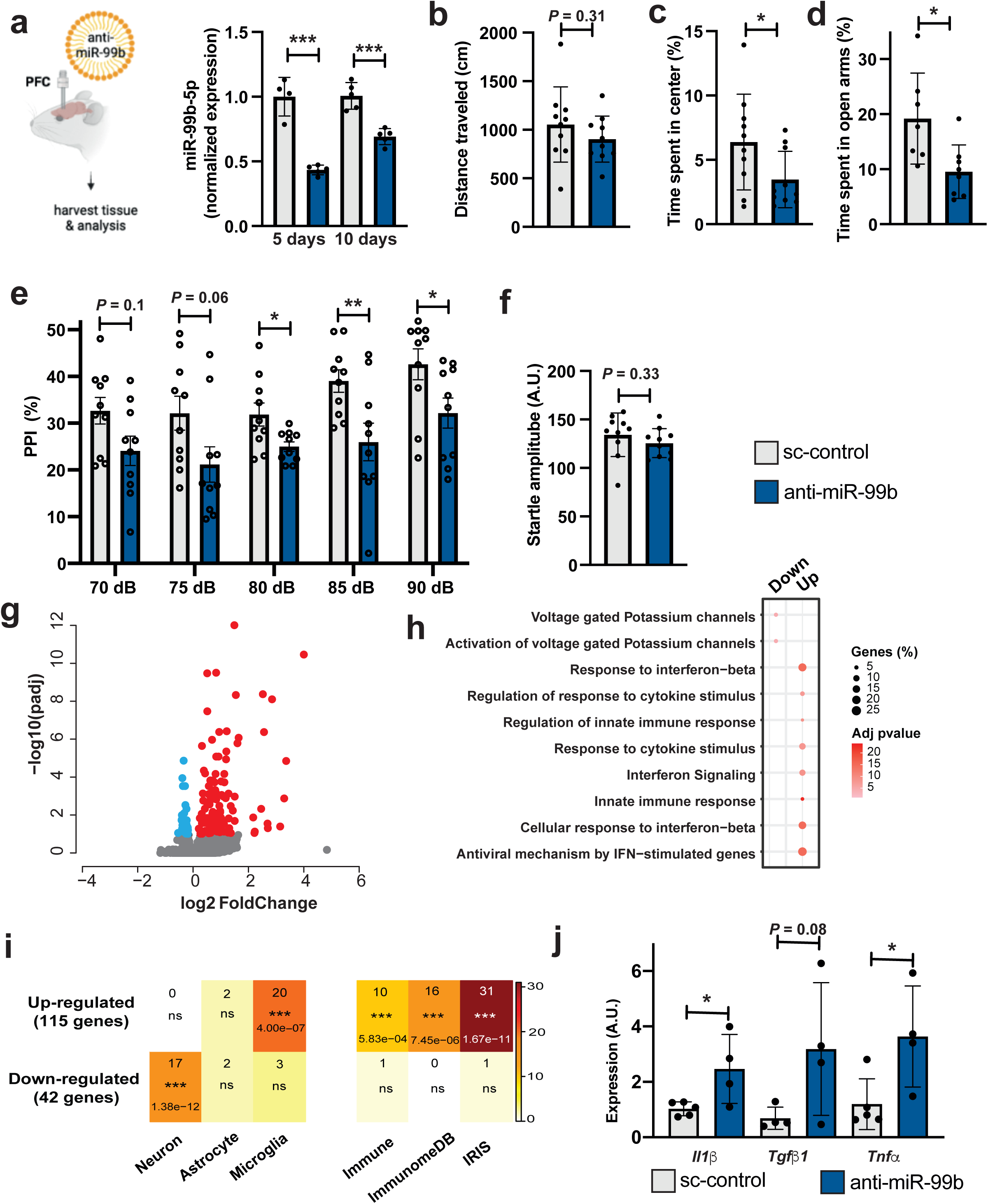
Decreasing miR-99b-5p levels in the PFC of mice leads to SZ-like phenotypes and increases the expression of genes linked to microglia activation. **a.** Left panel: Experimental design. Right panel: Bar graph showing qPCR results for miR-99b-5p in tissue obtained from the PCF of mice 5 or 10 days after injection of anti-miR-99b or sc-control oligonucleotides. (n=4/group; ****P < 0.0001, unpaired t test). **b**. Bar graph showing the distance traveled in the open field test of mice injected to the PFC with either anti-miR-99b or sc-control oligonucleotides (n=10/group; unpaired t test. **c.** Bar graph showing the time spent in the center of the open field in mice injected to the PFC with either anti-miR-99b or sc-control oligonucleotides (n=10/group; *P < 0.05; unpaired t test). **d.** Bar graph showing the time spent in the open arms when an elevated plus maze test was performed in mice injected to the PFC with either anti-miR-99b or sc-control oligonucleotides (n=10/group; *P < 0.05; unpaired t test). **e.** Bar graph showing the results of a PPI experiment of mice injected with either anti-miR-99b or sc-control oligonucleotides. PPI is impaired in anti-miR-99b injected mice (n=10/group) unpaired t test; *P < 0.05; **P < 0.01, ***P < 0.001; ****P < 0.0001). **f.** Bar graph showing the basic startle response among groups. **g.** Volcano plot showing the differentially expressed genes (upregulated in red, downregulated in blue) when RNA-seq was performed from the PFC of mice injected with either anti-miR-99b or sc-control oligonucleotides. Genes with log2-fold change ± 0.5 and adjusted p value < 0.05 are highlighted. **h.** GO-term analysis of the upregulated genes found in (e). **i.** Heat maps showing the enrichment of the upregulated genes as determined in (e) in various datasets. Left panel shows that the upregulated genes are enriched for microglia-specific genes, while the downregulated genes are enriched for neuron-specific genes. The right panel shows that the upregulated genes are over-represented in 3 different databases for immune function-related genes. **j.** Bar graph showing the qPCR results of the *Il1ß*, *Tgfb1* and *Tnfa* genes in FACS-sorted microglia collected from the PFC of mice injected with anti-miR-99b or sc-control oligonucleotides. (n=4 or 5/group; unpaired t test; *P < 0.05). Error bars indicate SD.

To gain first insights into the molecular processes controlled by miR-99b-5p in the brain, we injected another group of mice with anti-miR-99b and sc-control oligonucleotides and isolated PFC tissue 5 days later for RNA sequencing analysis. Differential expression analysis revealed 147 deregulated genes (adjusted *p*-value < 0.1, log2FC +/-0.2), of which 113 genes were upregulated and 34 were downregulated **(Fig. 2g; supplemental table 7)**. Gene ontology (GO) analysis revealed that upregulated genes were linked to processes such as innate immunity and interferon signaling **(Fig. 2h)**. GO analysis of the downregulated genes did not yield any highly significant pathways but detected processes linked to voltage-gated potassium channels **(Fig 2h, supplemental table 8)**. These data suggest that miR-99b-5p in the PFC may regulate mRNAs linked to immune-related processes. To further test this hypothesis, we compared the list of upregulated genes with gene expression data from neurons, astrocytes and microglia as well as genes present within 3 different immune function-related gene expression databases. We observed that the upregulated genes were highly enriched in microglia (*P* = 4 × 10^-12^), while no enrichment was observed in astrocytes or neurons. The downregulated genes were significantly enriched in neurons (*P* = 1.38 × 10^-12^). In line with this, the upregulated genes were significantly overrepresented in 3 databases for genes linked to immune function, namely the immune, immunome and IRIS databases **(Fig. 2i)**. Together, these data suggest that the levels of miR-99b-5p in the PFC of mice may specifically increase the expression of immune-related genes in microglia. To further test this we administered anti-miR-99b or sc-control to the PFC of mice and subsequently isolated CD45^low^/CD11b^+^ microglial cells via fluorescence-activated cell sorting (FACS). While microglia cell numbers did not differ between groups, miR-99b-5p expression was significantly decreased in microglia isolated from anti-miR-99b-treated mice **(Fig. EV4).** The expression of selected pro-inflammatory genes *Il1ß, Tgfb1* and *Tnfα* that were upregulated in the RNA-seq dataset were also increased in microglia obtained from anti-miR-99b-treated mice, although *Tgfβ1* failed to reach significance (*P* = 0.08) **(Fig 2j)**.

These data suggest that miR-99b-5p controls microglia-mediated immunity in the PFC, which is in agreement with previous studies linking aberrant microglia function and neuroinflammation to the pathogenesis of SZ [34].

### miR-99b-5p controls microglia-mediated immune function and affects dendritic spine number

To further explore the role of miR-99b-5p in microglia, we cultured primary microglia from the cortex of mice and treated these cells with anti-miR-99b or sc-control LNAs followed by RNA sequencing **(Fig 3a)**. Differential expression analysis revealed 139 deregulated genes, of which 104 were up- and 35 were downregulated **(Fig 3b, supplemental table 9)**. GO-term and KEGG-pathway analysis revealed that the upregulated genes were linked to immune activation and phagocytosis **(Fig 3c; supplemental table 10)**. In agreement with the *in vivo* data, we observed an increased expression of *Il1ß, Tgfb1, and Tnfα*, which could be confirmed via qPCR **(Fig 3d; see also Fig EV4c)**. Next, we investigated phagocytosis, a key function of microglia that is altered during neuroinflammation [35]. In a first approach we employed immortalized microglia (IMG) cells. Similar to the treatment with the lipopolysaccharide (LPS) commonly used to induce microglia activation, inhibition of miR-99b-5p caused an upregulation of proinflammatory cytokines and increased phagocytosis as measured via the uptake of fluorescent latex beads **(Fig. EV5)**. Encouraged by these data we performed similar experiments in primary microglia isolated from the mouse PFC. In line with the data obtained in IMG cells, treatment of primary microglia with anti-miR-99b LNAs significantly increased phagocytosis **(Fig 3e)**.

**Figure 3:**
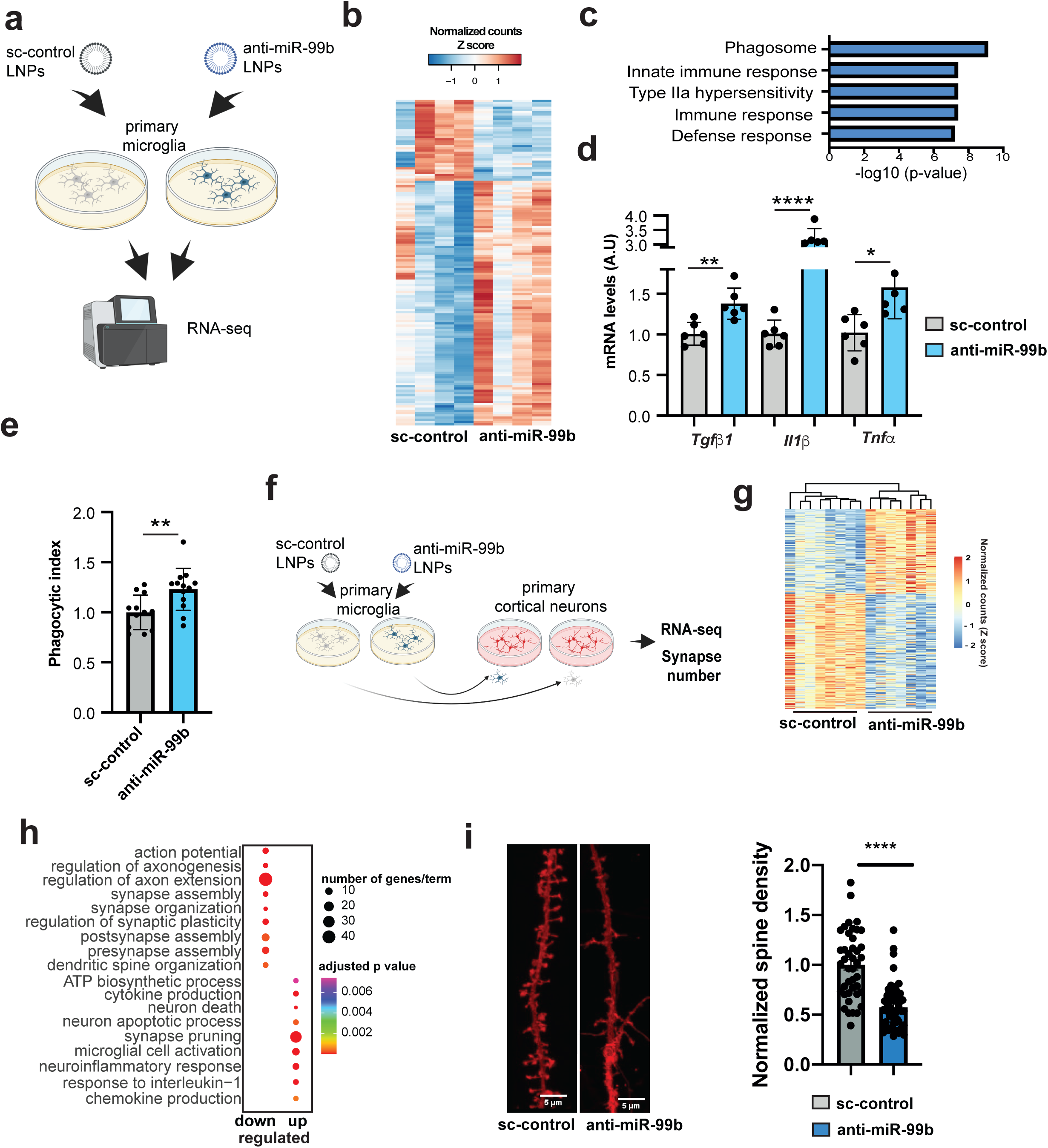
Decreasing miR-99b-5p levels in microglia increases phagocytosis and reduces synapse number in cortical neurons. **a.** Left panel: Experimental design. **b.** Volcano plot showing differential expressed genes when comparing microglia treated with anti-miR-99b or sc-control LNAs. Genes with statistical significance are highlighted. **c.** Bar chart showing the top GO terms represented by the up regulated genes shown in (b). **d.** Bar charts showing qPCR results for *Tgfb1*, *Il1ß* and *Tnfa* comparing microglia treated with anti-miR-99b or sc-control LNAs (n=6/group; unpaired t test; *P < 0.05; **P < 0.01, ****P < 0.0001). **e.** Bar chart showing the results of a phagocytosis assay performed in microglia treated with anti-miR-99b in comparison to cells treated with sc-control LNAs. The percentage of phagocytic index represents (# of total engulfed beads in an image / # of total cells identified in an image; n = 13 independent experiments; unpaired t test; **P < 0.01). **f**. Experimental scheme illustrating the co-culture experiment. **g.** Heat map showing the differentially expressed genes from the experiment described in (f). **h.** Plot showing the results of a GO term analysis for the up- and downregulated genes displayed in (g). **i.** Left panel: Representative image showing DIL dye staining to visualize dendritic spines in co-cultures as illustrated in (f). Scale bar 5 μm. Right panel: Bar chart showing the statistical quantification of the data depicted in (i). Each dot represents a spine density for a dendritic segment. *P < 0.05; **P < 0.01, ***P < 0.001; ****P < 0.0001). Error bars indicate SD.

Aberrant microglia activity can have detrimental effects on neuronal plasticity [36]. To test whether the reduced expression of miR-99b-5p in microglia could affect neuronal plasticity, we performed a co-culturing experiment. Primary microglia were first treated with sc-control or anti-miR-99b LNAs for 48 h before being harvested and transferred to cortical neuronal cultures **(Fig 3f)**. RNA was isolated from these co-cultures after 48 h and subjected to RNA-seq. Differential expression analysis revealed 366 deregulated genes (155 upregulated and 211 downregulated genes, adjusted p value < 0.05, log2FC +/-0.1); **Fig. 3g, supplemental table 11**). GO term analysis of the upregulated genes revealed neuroinflammatory processes, neuron death, neuron apoptotic processes as well as synaptic pruning **(Fig 3h, supplemental table 12_UP)**, while downregulated genes were associated with processes indicating loss of synaptic function such as regulation of axon extension or synapse organization **(Fig 3h, supplemental table 12_DOWN)**.

These data are in agreement with our previous findings suggesting that loss of miR-99b-5p increases inflammatory processes in microglia. More importantly, the data suggest that microglia lacking miR-99b-5p may have detrimental effects on synaptic function when co-cultured with cortical neurons. It is particularly interesting that synaptic pruning is detected as a major process increased in the co-cultures, since synaptic pruning has been linked to SZ [36]. In line with this, key factors of the complement system known to drive pathological synaptic pruning were increased in primary microglia treated with anti-miR-99b, as well as in the corresponding co-cultures and also in the postmortem human prefrontal cortex of schizophrenia patients **(Fig EV6)**. When we analyzed the number of dendritic spines, we observed that spine density was significantly reduced in neurons co-cultured with microglia that had received anti-miR-99b, when compared to cultures treated with corresponding control microglia **(Fig 3 i)**.

### miR99b-5p control neuroinflammation via the regulation of Zbp1

The three RNAseq datasets obtained from the PFC of mice, primary microglia and the co-cultures consistently show that knockdown of miR-99b-5p increases the expression of genes linked to inflammatory processes. Many of the gene expression changes likely represent secondary effects. To better understand the mechanisms by which miR-99b-5p controls neuroinflammation, we aimed to identify direct targets of miR-99b-5p (**supplemental table 13**). When we analyzed the RNA-seq data obtained from the PFC (see Fig 2), we identified 13 out of 113 genes as potential mRNA targets of miR99b-5p **(Fig 4A)**. GO term analysis was performed for the 13 genes and revealed that they are linked to inflammatory processes including type I interferon signaling **(Fig. EV7, supplemental table 14)**, linked to schizophrenia [37] [38]. Seven of these genes were also upregulated in primary microglia treated with anti-miR-99b, and among them were key regulators of inflammatory processes such as *Stat1* which was found to be hyperactive in blood samples of SZ patients [39]. A gene that specifically caught our attention was *Zbp1* because the corresponding protein - also known as the DNA-dependent activator of interferon regulatory factors (*Dai*) - is a key regulator of pro-inflammatory processes that result in the activation of inflammatory caspases and the induction of *Il1ß* [27]. Thus, *Zbp1* represented a rather upstream factor in the inflammatory cascade. On this basis, we speculated that the regulation of *Zbp1* could be a key mechanism by which miR-99b-5p regulates inflammatory processes and may contribute to the pathogenesis of SZ when it is increased.

**Figure 4:**
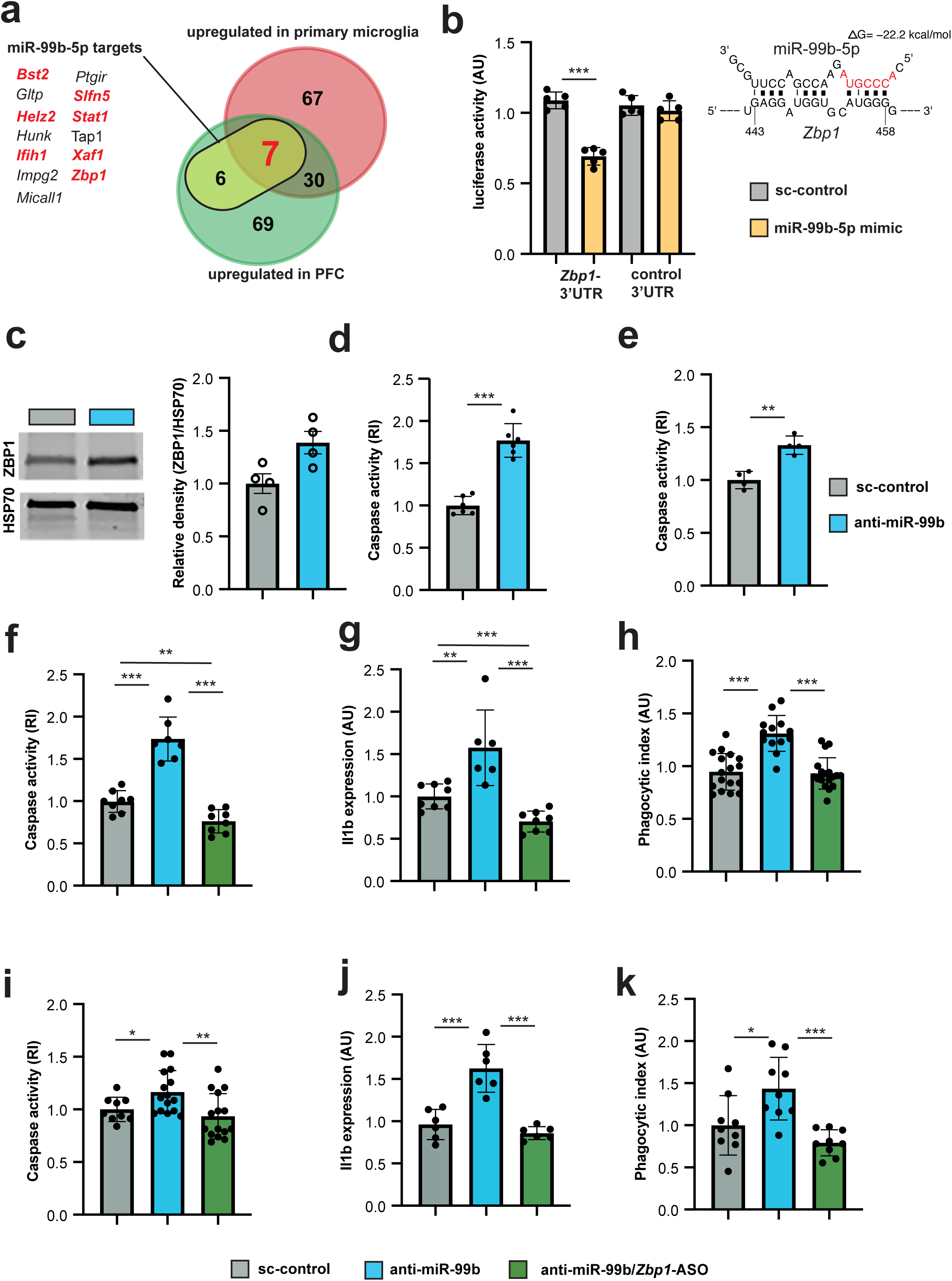
miR-99b-5p regulates neuroinflammatory phenotypes via *Zbp1*. **a.** Venn diagram comparing the genes upregulated in the PFC of mice and in primary microglia when injected or treated with anti-miR99b vs. sc-control LNAs, respectively. The data is further compared to the identified 13 miR-99b-5p target mRNAs detected in the PFC dataset. The left panel shows the gene names of the 13 miR-99b-5p target mRNAs. Red indicates miR-99b-5p targets upregulated in the PCF and in primary microglia upon anti-miR-99b treatment. **b.** Bar graph showing the results of the luciferase assay. In comparison to sc-control LNAs, administration of miR-99b-5p mimic decreases luciferase activity when cells express the Zbp1-3’UTR. This effect is not observed when a control 3’UTR that does not bind miR-99b-5p is used. (n=6/group). The upper right panel shows the predicted binding of miR-99b-5p to the 3’UTR of *Zbp1*. **c.** Left panel: Representative immunoblot image showing ZBP1 levels in microglia treated with sc-control LNAs or anti-miR-99b. HSP70 was used as a loading control. Right panel: Bar graph showing the quantification of the data depicted in the left panel. n=4/group. **d.** Bar graph showing quantification of caspase activity in primary microglia treated with sc-control LNAs or anti-miR-99b (n=6/group). **e.** Bar graph showing quantification of caspase activity in protein lysates isolated from the PFC of mice injected with anti-miR-99b or sc-control (n=4/group). **f.** Bar graph showing quantification of caspase activity in primary microglia treated with either sc-control LNAs, anti-miR-99b or anti-miR-99b together with *Zbp1*-ASOs. (n=6/group). **g.** Bar graph showing qPCR results for *Il1ß* in primary microglia treated with either sc-control LNAs, anti-miR-99b or anti-miR-99b together with *Zbp1*-ASOs (n=6/group). **h.** Bar graph showing the results of a phagocytosis assay performed in primary microglia treated with either sc-control LNAs, anti-miR-99b or anti-miR-99b together with *Zbp1*-ASOs (n=16 independent experiments). **i.** Bar graph showing quantification of caspase activity in human iPSC-derived microglia treated with either sc-control LNAs, anti-miR-99b or anti-miR-99b together with *Zbp1*-ASOs (n=13-16 samples/group). **j.** Bar graph showing qPCR results for IL1ß in human iPSC-derived microglia treated with either sc-control LNAs, anti-miR-99b or anti-miR-99b together with *Zbp1*-ASOs (n=6/group). **k.** Bar graph showing the results of a phagocytosis assay performed in human iPSC-derived microglia treated with either sc-control LNAs, anti-miR-99b or anti-miR-99b together with *Zbp1*-ASOs. The percentage of phagocytic index represent (# of total engulfed beads in an image / # of total cells identified in an image; n = 9 independent experiments. Error bars indicate SD; unpaired t test; *P < 0.05; **P < 0.01, ***P < 0.001; ****P < 0.0001). RI: relative immunofluorescent,

First, we performed a luciferase assay to directly test the regulation of *Zbp1* by miR-99b-5p. We used the renilla dual luciferase reporter vector harboring the *Zbp1*-3ʹUTR. Co-transfection of this vector with miR-99b-5p LNAs significantly reduced the luciferase activity **(Fig 4b)**, but this was not the case when scramble control LNAs were used **(Fig 4b)**. Moreover, we observed that ZBP1 protein levels were significantly increased in primary microglia treated with anti-miR-99b **(Fig 4c)**. These data show that miR-99b-5p can directly regulate *Zbp1* levels.

On this basis we decided to investigate whether the inflammatory phenotypes induced in response to decreased miR-99b-5p levels depend on *Zbp1*. So far, we have found that decreased levels of miR-99b-5p lead to enhanced phagocytosis and increased expression of pro-inflammatory cytokines such as *Il1ß* which has been associated with *ZBP1* activity [40]. Another important step in ZBP1-mediated orchestration of inflammation is the activation of pro-inflammatory caspases [41] [42], and therefore we examined whether reduced miR-99b-5p levels would also affect the activity of pro-inflammatory caspases. Indeed, when primary microglia were treated with anti-miR-99b, caspase activity was significantly increased when compared to that in cells treated with sc-control LNAs **(Fig. 4d)**. Similar findings were obtained when protein lysates isolated from the PFC of mice injected with either anti-miR-99b or sc-control LNAs were analyzed for caspase activity **(Fig. 4e)**.

To test whether the miR-99b-5p-mediated increase in caspase activity, *IL-1ß* expression and phagocytosis depends on *Zbp1*, we treated primary microglia with either anti-miR-99b alone or in combination with an anti-sense oligonucleotide (ASO) targeting *Zbp1* (*Zbp1*-ASO). In agreement with our previous observation, anti-miR-99b treatment increased caspase activity. This effect was ameliorated in microglia treated with anti-miR-99b and *Zbp1*-ASO **(Fig 4f)**. Similar observations were made when we analyzed *IL1ß* expression **(Fig. 4g)** and phagocytosis **(Fig 4h)**. These data suggest that ZBP-1 plays an important role in mediating the neuroinflammatory processes downstream of miR-99b-5p.

We performed parallel experiments in human iPSC-derived microglia. Similar to the mouse data, administration of anti-miR-99b increased caspase activity **(Fig 4i)**, IL-1ß expression **(Fig 4j)** and phagocytosis **(Fig 4k)** when compared to human iPSC-derived microglia treated with sc-control LNAs. These effects were attenuated when anti-miR-99 LNAs were co-administered with *Zbp1*-ASOs **(Fig 4i-k)**. These data suggest that in human microglia, miR-99b-5p also controls neuroinflammatory processes via the regulation of *Zbp-1* expression. In line with this interpretation, IL1ß and Zbp1 mRNA levels were increased in postmortem human brain samples from SZ patients and controls **(Fig. S8a)**. To determine whether knockdown of *Zbp1* would also mitigate the effect of anti-miR-99b treatment on SZ-like behavior in mice, we injected either anti-miR-99b alone or in combination with *Zbp1*-ASOs into the PFC of mice before subjecting the animals to behavior testing. Mice injected with sc-control LNAs served as controls. The corresponding data revealed that *Zbp1* knockdown rescues anti-miR-99b-mediated impairment of PPI **(Fig. S8)**.

Our findings suggest that reduced miR-99b-5p levels in microglia contribute to the pathogenesis of schizophrenia via the regulation of *Zbp1*-controlled neuroinflammation. Therefore, miR-99b-5p may constitute a novel biomarker for SZ, while targeting miR-99b-5p and/or ZBP1 might represent an effective SZ treatment

## Discussion

In this study we combined the analysis of blood samples and postmortem brain tissue to identify miRs involved in the pathogenesis of SZ. Using WGCNA as well as differential expression analysis in blood samples, we identified several miRs that differed between patients and controls and were significantly correlated with SZ phenotypes. GO term analysis of the confirmed target genes of these miRs hinted at a number of molecular processes of which pathways linked to immune function were overrepresented. Such a GO term analysis based on miR target genes is, of course, not ultimately conclusive but our observation is in agreement with previous studies showing that neuroinflammation plays a role in the pathogenesis of SZ [43] [44]. To further refine the identification of miRs linked to SZ we also performed a differential expression analysis of the small RNA seq data obtained from blood as well as from postmortem brain tissue of SZ patients and controls. We eventually identified five candidate miRs, miR-101-3p, miR-378a-3p, miR-21-5p, miR-192-5p and miR-103a-3p, that were increased in the blood and brain of SZ patients. All of these miRs have been implicated brain and non-brain diseases. For example, knock-down of miR-101-3p in the hippocampus impairs learning in mice [45] while increased circulating levels of miR-101-3p have been observed in patients suffering from autism [46], attention-deficit hyperactivity disorder [47] but also diabetes [48] [49] or cancer [50]. MiR-378a-3p has been associated with cerebral ischemia [51] is increased in blood samples of Down syndrome patients [52] and was reported to be part of a blood-miR signature that can distinguish Alzheimer’s disease patients from control [53]. miR-21-5p was found to be increased in blood samples of SZ patients [29], while antipsychotic treatment is correlated with decreased miR-21-3p expression [54]. Moreover, miR-21-3p was altered in blood samples of patients suffering from bipolar disease [55]. However, altered miR-21-3p has also been observed in various other diseases and is for example increased in patients with lung cancer [56]. Several studies have linked miR-192-5p to cognitive function and depression. For example, miR-192-5p was decreased in the blood AD patients upon aerobic exercise [57] and was altered in the brains of patients suffering from major depressive disorder [58], while increasing the levels of miR-192-5p in a mouse model for depression ameliorated cognitive impairments [59]. Finally, altered blood levels of miR-103-3p have been linked to childhood traumatization and depression [60] as well as Autism [61].

Four miRs, miR-500a-3p, miR-501-3p, miR-221-5p and miR-99b-5p, that all originated from the ME_Turquoise co-expression module which was downregulated in SZ patients and significantly correlated to SZ phenotypes, were also decreased in the postmortem brains of SZ patients . A recent study identified miR-501-3p as a schizophrenia-associated miR as it was found to be decreased in blood samples of monozygotic twins discordant for SZ [21]. The authors went on to show that loss of miR-501-3p in mice leads to SZ-like phenotypes, a finding that was linked to miR-501-3p-mediated regulation of *metabotropic glutamate receptor 5* expression in the cortex. These data are in agreement with our observation that miR-501-3p was decreased in SZ patients of the PsyCourse study as well as in the postmortem brains of SZ patients. As to the other 3 miRs, there are thus far no data on the role of miR-500a-3p in the CNS, and while miR-221-5p has recently been linked to the regulation of synaptic processes [62], there are still also no data on the role of miR-221-5p in SZ. As regards to miR-99b-5p, no functional data are available on its role in the CNS, and no report has as yet implicated this miR in the pathogenesis of SZ.

In summary, we have discovered miRs that have already been linked to the pathogenesis of SZ or other brain diseases, as well as miRs that have not been extensively studied so far. These data suggests that our approach may provide a viable means to detect novel SZ-associated miRs.

To test this hypothesis, we decided to investigate miR-99b-5p that was part of the ME_Turquoise co-expression module but was also decreased in the postmortem brain of SZ patients as well as in the blood of SZ patients, when the data was analyzed via differential expression. Moreover, essentially nothing was known about the role of miR-99b-5p in the adult brain. Inhibiting miR-99b-5p in the prefrontal cortex of mice led to impaired PPI and increased anxiety. PPI is impaired in SZ patients and in mouse models for SZ [32] [33]. Furthermore, increased anxiety is a phenotype often observed in SZ patients [30], suggesting that reduced miR-99b-5p levels are indeed linked to the development of SZ-like phenotypes. Nevertheless, these data cannot establish a clearly causal link between reduced miR-99b-5p expression and the pathogenesis of SZ in humans, since no animal model can fully recapitulate the complex processes in human patients due to functional and structural differences in cortical anatomy [63] [64].

However, that miR-99b-5p is involved in the pathogenesis in SZ is further underscored by the results of our molecular analysis. RNA-seq analysis of the prefrontal cortex of mice revealed that inhibition of miR-99b-5p mainly led to an increased expression of genes, which is in agreement with the established action of miRs in controlling mRNA levels. Furthermore, the upregulated genes were almost exclusively related to immunity pathways in microglia, a process which has been linked to the pathogenesis of SZ by various means. For example, altered microglia have been observed in postmortem brain samples of SZ patients [65]. In addition, epidemiological data demonstrated a correlation between immune diseases and SZ [66], while several neuroimaging studies reported an increase in activated microglia in the brains of SZ patients [67] [68]. Finally, studies in animal models have implicated aberrant microglia activation with the onset of SZ-like phenotypes [69] [70]. While miR-99b-5p has not been studied in microglia so far, these data are in line with previous reports demonstrating a role of the miR-99b in the modulation of inflammatory responses. For example, miR-99b levels are decreased in tumor-associated macrophages and re-expression of miR-99b attenuates tumor growth [71]. Furthermore, inhibition of miR-99b in dendritic cells significantly elevated the levels of proinflammatory cytokines including *Il1ß* and *Tnfα* [72]. These findings are in agreement with our data showing that inhibition of miR-99b-5p in the prefrontal cortex of mice increased the expression of pro-inflammatory cytokines including *Il1ß* and *Tnfα* in microglia that we had isolated from the brains of these mice via FACS. A strong upregulation of genes linked to inflammatory processes, including the upregulation of *Il1ß* and *Tnfα,* was also observed when miR-99b-5p was inhibited in IMG cells or primary microglia. This is interesting since increased *Ill1 ß* and *Tnfα* levels have been repeatedly reported in SZ patients [73] and may offer novel therapeutic avenues. For example, inhibition of TNF*α* was recently shown to ameliorate disease phenotypes in different mouse models of SZ [70].

Aberrant microglia activation can affect neuronal function via synaptic pruning, a process that is based on the phagocytic activity of microglia [74]. We observed that inhibition of miR-99b-5p in IMG cells and in primary microglia increased their phagocytic activity. Moreover, cortical neurons co-cultured with microglia that were treated with anti-miR-99b oligonucleotides displayed differentially expressed genes, of which the downregulated genes were linked to GO terms such as synapse assembly, regulation of synaptic plasticity or dendritic spine organization. As for the upregulated genes, the most significant GO term was synapse pruning. Since our data also revealed that neurons co-cultured with anti-miR-99b-treated microglia indeed displayed a reduced number of dendritic spines, our findings suggest a scenario in which reduced levels of miR-99b-5p lead to an upregulation of proinflammatory processes in microglia, which eventually impacts on synaptic structure. This interpretation is in agreement with previous reports suggesting that aberrant microglia activation leads to pathological synaptic pruning, which in turn leads to plasticity defects which could drive the pathogenesis of SZ [36] [75]. Notably, the increased expression of several complement factors in microglia have been implicated in this process [76]. In line with these data, we observed increased expression of key complement factors in primary microglia and in corresponding microglia/neuron co-cultures in which miR-99b-5p was inhibited, as well as in postmortem brain samples from SZ patients. In summary, these findings provide a plausible mechanism on how reduced levels of miR-99b-5p can contribute to the pathogenesis of SZ, namely the induction of a pro-inflammatory response associated with synaptic pruning. Nevertheless, we cannot exclude that additional mechanisms within microglia or other neural cells play a role.

MiRs mediate their biological action by controlling the expression of specific target mRNAs. Our data showed that within microglia, miR-99b-5p controls the expression of the *Zbp1* gene that plays an important role in the innate immune response [27]. ZBP1 acts as sensor for Z-DNA/Z-RNA and controls inflammatory pathways such as type I interferon-signaling and other pathways, eventually leading to the upregulation of various pro-inflammatory cytokines including e.g. the induction of *IL1ß* [77] [78] [40].

These data suggest that reduced levels of miR-99b-5p in microglia contribute to SZ-like phenotypes because the tight control of *Zbp1* levels is lost. In line with this interpretation, we demonstrated that the administration of *Zbp1*-ASO rescues the effects of anti-miR-99b treatment on SZ-like phenotypes in mice as well as in the corresponding cellular alterations observed in primary microglia from mice as well as in microglia derived from human iPSCs. Interestingly, the cellular processes we find to be affected by altered miR-99b-5p and *Zbp1* levels have also been implicated in other brain diseases. Thus, it will be important to investigate the role of miR-99b-5p and *Zbp1* in other neuropsychiatric diseases. Moreover, aberrant microglia activation and synaptic pruning is observed in neurodegenerative diseases such as Alzheimer’s disease [79], and ZBP1 also controls the NRLP3 inflammasome [78], a key regulator of neuroinflammatory phenotypes in Alzheimer’s disease [80]. In this context it is interesting to note that one study found decreased miR-99b-5p levels in plasma samples obtained in a mouse model of Alzheimer’s disease when measured at 6 and 9 months of age, while increased levels were reported in older mice [81]. These data might underscore the need for further study as to the role of *miR-99b-5p* and *Zbp1* in microglia obtained from wild type mice as well as mouse models for neuropsychiatric or neurodegenerative diseases at different ages. Indeed, it is well established that microglia undergo age-dependent functional changes and even differ between brain regions [82] [83].

There are other questions we could not address within the scope of this manuscript. It will for example be interesting to investigate the other candidate miRs we found in addition to miR-99b-5p. Similarly, it will be important to study the potential miR-99b-5p targets we found in addition to *Zbp-1* in the context of SZ. Another question relates to the mechanisms that underlie the downregulation of miR-99b-5p in SZ patients. In future projects it will be interesting to test for example whether miR-99b-5p is altered in SZ mouse models that are based on either genetic or environmental risk factors such as early life stress. In addition, it will be important to identify the source of elevated miR-99b-5p levels in blood samples of SZ patients. It is known that miRs can be transported from the brain to the periphery within exosomes [84] [25], and recent studies reported the isolation of microglia-derived exosomes from human blood [85]. While this approach is not undisputed, it will be interesting to apply such methods to the PsyCourse Study, which is, however, beyond the scope of the current work. Although our findings that miR-99b-5p is decreased in the brain and the blood of SZ patients support the idea that the changes in blood may reflect corresponding changes in the brain, we cannot conclusively answer this question at present. Rather, we suggest that the analysis of miR-99b-5p levels in blood may eventually help stratify patients for treatment, including novel approaches based on RNA therapeutics towards miR-99b-5p or *Zbp1*.

In conclusion, in the present study we identify a miR-99b-5p-*Zbp1* pathway in microglia as a novel mechanism that likely contributes to the pathogenesis of schizophrenia. Our data also suggest that strategies to increase the levels of miR-99b-5p or inhibit *Zbp1*, for example via ASOs, could serve as novel therapeutic strategies for treating SZ patients.

## Material and Methods

### Human subjects

All experiments involving human data were approved by the relevant Ethics committees (see Budde et al.). Informed written consent was obtained for all subjects. Blood samples (PAXgene Blood RNA Tubes; PreAnalytix, Qiagen) and behavioral data (supplemental table 1) of control and schizophrenia patients were obtained from participants of the PsyCourse Study [26]. Psychiatric diagnoses were confirmed using the Diagnosis and Statistical Manual of Mental Disorders Fourth Edition (DSM-IV) criteria. Control subjects were screened for psychiatric disorders using parts of the structured clinical interviews for mental disorders across the lifespan (MINI-DIPS). All subjects were assessed for psychiatric symptoms through a battery of standard tests including the Positive and Negative Syndrome Scale (PANSS), the Global Assessment of Functioning Scale (GAF), and the Beck depression inventory (BDI-II).

### Post-mortem human brain samples

Postmortem tissue samples (prefrontal cortex A9&24) from controls (n = 17; 5 females & 12 males; age = 62.3 ± 18.9 years, PMD = 19.7 ± 6.7 h) and schizophrenia patients (n = 13; 5 females & 8 males; age 57.7 ± 16.8 years, PMD = 21 ± 6.4 h) were obtained with ethical approval and upon informed consent from the Harvard Brain Tissue Resource Center (Boston, USA). RNA was isolated using Trizol as described in the manufacturer protocol using the Directzol RNA isolation kit (Zymo Research, Germany). RNA concentration was determined by UV measurement. RNA integrity for library preparation was assessed using an RNA 6000 NanoChip in a 2100 Bioanalyzer (Agilent Technologies).

### High throughput small RNAome sequencing

Small RNAome libraries were prepared with total RNA according to the manufacturer’s protocol with NEBNext® small RNA library preparation kit. All human subject small RNAome libraries were prepared with 150 ng of total RNA. Briefly, total RNA was used as starting material, and the first strand of cDNA was generated, followed by PCR amplification. Libraries were pooled and PAGE was run for size selection. For small RNAome, ∼150 bp band was cut and used for library purification and quantification. A final library concentration of 2 nM was applied for sequencing. The Illumina HiSeq 2000 platform was used for sequencing and was performed using a 50-bp single read setup. Illumina’s conversion software bcl2fastq (v2.20.2) was used for adapter trimming and converting the base calls in the per-cycle BCL files to the per-read FASTQ format from raw images. Demultiplexing was carried out using Illumina CASAVA 1.8. Sequencing adapters were removed using cutadapt-1.8.1. Sequence data quality was evaluated using FastQC (http://www.bioinformatics.babraham. ac.uk/projects/fastqc/). Sequencing quality was determined by the total number of reads, the percentage of GC content, the N content per base, sequence length distribution, duplication levels, overrepresented sequences and Kmer content.

### Data processing, QC, and Differential expression (DE) analysis

Sequencing data was processed using a customized in-house software pipeline. Quality control of raw sequencing data was performed by using FastQC (v0.11.5). The quality of miRNAs reads was evaluated by mirtrace (v1.0.1). Reads counts were generated using TEsmall (v0.4.0) which uses bowtie (v1.1.2) for mapping. Reads were aligned to the Homo_sapiens GRCh38.p10 genome assembly (hg38). The miRNA reads were annotated using miRBase. Read counts were normalized with the DESeq2 (v1.26.0) package. Unwanted variance such as batch effects, library preparation effects, or technical variance was removed using RUVSeq for all data (v1.20.0; k = 1 was used for factors of unwanted variation). DeSeq2 was utilized for differential expression analysis and adjustment of confounding factors. In the DESeq2 model, the PsyCourse data were corrected for sex, age and medication in DeSeq2. Volcano plots were plotted with the R package EnhancedVolcano (v1.4.0).

### WGCNA analysis

microRNAome co-expression module analysis was carried out using the weighted gene co-expression network analysis (WGCNA) package (version 1.61) in R [86]. We first regressed out age, gender, and other latent factors from the sequencing data, and after that, normalized counts were log (base 2) transformed. Next, the transformed data were used to calculate pairwise Pearson’s correlations between microRNAs and define a co-expression similarity matrix, which was further transformed into an adjacency matrix. Next, a soft thresholding power of 8 was chosen based on approximate scale-free topology and used to calculate pairwise topological overlap between microRNAs in order to construct a signed microRNA network. Modules of co-expressed microRNAs with a minimum module size of 10 were later identified using cutreeDynamic function with the following parameters: method = “hybrid”, deepSplit =4, pamRespectsDendro =F, pamStage = T. Closely related modules were merged using a dissimilarity correlation threshold of 0.25. Different modules were summarized as a network of modular eigengenes (MEs), which were then correlated with the different psychiatric symptoms and functionality variables (e.g., PANSS, GAF etc). The module membership (MM) of microRNAs was defined as the correlation of microRNA expression profile with MEs, and a correlation coefficient cutoff of 0.5 was set to select the module specific microRNAs. The Pearson correlation of MEs and psychiatric symptoms and functionality variable was plotted as a heat map.

### Enriched gene ontology and pathways analysis

To construct the Gene Regulatory network (GRN) for miRNA-target genes we retrieved validated microRNA targets from miRTarBase (v 7.0) (http://mirtarbase.mbc.nctu.edu.tw/). microRNA target genes were further filtered based on the expression in the brain. Brain-enriched expression was examined using the Genotype-Tissue Expression (GTEx) database. (GTEx Consortium). To identify the biological processes and their pathways in the miRNA-target genes, the ClueGO v2.2.5 plugin of Cytoscape 3.2.1 was used [87]. In the ClueGo plugin [88] a two-sided hypergeometric test was used to calculate the importance of each term and the Benjamini-Hochberg procedure was applied for the P value correction. KEGG (https://www.genome.jp/kegg/) and Reactome (https://reactome.org/) databases were used for the pathway analysis. To construct GRN for significantly deregulated mRNAs, the ClueGO v2.2.5 plugin of Cytoscape 3.2.1 was used. Biological processes (BP) and pathways with adjusted p value < 0.05 were selected for further analysis. For further analysis, cellular metabolism and cancer-related biological processes were omitted. Key BPs with low levels of GOLevel (because terms at lower levels are more specific and terms higher up are more general) were further considered for data presentation and interpretation.

### microRNA and mRNAs lipid nanoparticles preparation

miR99b-5p inhibitor sequences were used to decrease the expression of miR99b-5p. To decrease the expression of mRNAs, anti-sense oligos (ASO) were employed. ASOs, inhibitor and negative control sequences were purchased from Qiagen. MicroRNA inhibitor, or ASOs lipid nanoparticle (LNP) formulation, was achieved using a proprietary mixture of lipids containing an ionizable cationic lipid, supplied as Neuro9™ siRNA Spark™ Kit (5 nmol). The miRNA inhibitor or ASOs were encapsulated using a microfluidic system for controlled mixing conditions on the NanoAssemblr™ Spark™ system (Precision Nanosystems, Canada). The experiments were performed as described in the manufacturer’s protocol. Briefly, 5 nmol lyophilized microRNA inhibitor or ASOs were dissolved in formulation buffer 1 (FB1) to a final concentration of 2 nmol. This solution was further diluted to a final concentration of 930 ug/mL. Formulation buffer 2 (FB 2), microRNA inhibitor/ASOs in FB1, and lipid nanoparticles were added to the cartridge and encapsulated using the NanoAssembler Spark system.

### Animals

C57BL/6J mice were purchased from Janvier and housed in an animal facility with a 12-h light–dark cycle at constant temperature (23 °C) with *ad libitum* access to food and water. Animal experiments complied with relevant ethical regulations and were performed as approved by the local ethics committee. All experiments were performed with 3 months old male mice. Pre-frontal cortex (PFC) region was dissected on day five after stereotaxic surgery for RNA-seq-based experiments.

### Stereotaxic surgery

For intracerebral stereotaxic injections of LNPs in the PFC, 3-month-old mice were anesthetized with Rompun 5mg/kg and Ketavet 100mg/kg. After application of local anesthesia to the skull, two small holes were drilled into the skull. Mice then received a bilateral injection of LNPs of microRNA inhibitor/negative control or ASOs (dose: 0.15 ug/mL for microRNA inhibitor/negative control; dose: 0.3 ug/mL for ASO+ microRNA inhibitor mix). LNPs were injected with a rate of 0.3 μl/min per side. Only 0.9 ul of LNPs were injected per hemisphere (0,5 µl/min). After surgery, all mice were monitored until full recovery from the anesthesia and housed under standardized conditions.

### Behavioral phenotyping

The open field test was performed to evaluate locomotory and exploratory functions. Mice were placed individually in the center of an open arena (of 1 m length, 1 m width, and side walls 20 cm high). Locomotory activity was recorded for 5 min using the VideoMot2 tracking system (TSE Systems). The elevated plus maze test was used to evaluate basal anxiety. Mice were placed individually in the center of a plastic box consisting of two open and two walled closed arms (10 × 40 cm each, walls 40 cm high). Their behavior was recorded for 5 min using the VideoMot2 system. Time spent in open versus closed arms was measured to assay basal anxiety phenotype. Prepulse inhibition (PPI) was performed to test the acoustical startle response (ASR). ASR was completed in an enclosed sound-attenuated startle box from TSA Systems. In brief, mice were placed individually inside a cage attached with a piezoelectric transducer platform in a sound-attenuated startle cabinet. These sensory transducers converted the movement of the platform induced by a startle response into a voltage signal. Acoustic stimuli were executed through speakers inside the box. The mice were given 3 min to habituate at 65 dB background noise and their activity was recorded for 2 min as baseline. After the baseline activity recording, the mice were tested to six pulse-alone trials, at 120-dB startle stimuli intensity for a duration of 40 ms. PPI of startle activity was measured by conducting trials for pre-pulse at 120 dB for 40 ms or preceding non-startling prepulses of 70, 75, 80, 85, 90 dB.

### RNA isolation

#### Humans

PAXgene Blood RNA Tubes (PreAnalytix/Qiagen) were stored at -80°C. For RNA isolation, the tubes were thawed and incubated at room temperature overnight. RNA was extracted according to the manufacturer’s protocol using PAXgene Blood RNA Kits (Qiagen). RNA concentrations were measured by UV measurement. RNA integrity for library preparation was determined by analyzing them on an RNA 6000 NanoChip using a 2100 Bioanalyzer (Agilent Technologies).

#### Mice

The mice were sacrificed by cervical dislocation on day five after stereotaxic surgery. Unilateral PFC region was collected and immediately frozen in liquid nitrogen and later stored at -80°C until RNA isolation. Total RNA was isolated using the trizol method as described by the manufacturer’s protocol using the Directzol RNA isolation kit (Zymo Research, Germany). The RNA concentration was determined by UV measurement. RNA integrity for library preparation was assessed using a Bioanalyzer (Agilent Technologies).

### RNA sequencing

Total RNA was used for the library preparation using the TrueSeq RNA library prep kit v2 (Illumina, USA) according to the manufacturer’s protocol. 500 ng RNA was used as starting material. The quality of the libraries was assessed using the Bioanalyzer (Agilent Technologies). Library concentration was measured by Qubit™ dsDNA HS Assay Kit (Thermo Fisher Scientific, USA). Multiplexed libraries were directly loaded onto a Hiseq2000 (Ilumina) with 50 bp single read setup.

The sequencing data were processed using a customized in-house software pipeline. Illumina’s conversion software bcl2fastq (v2.20.2) was employed for adapter trimming and converting the base calls in the per-cycle BCL files to the per-read FASTQ format from raw images. Quality control of raw sequencing data was carried out using FastQC (v0.11.5)(http://www.bioinformatics.babraham. ac.uk/projects/fastqc/). Reads were aligned using the STAR aligner (v2.5.2b) and read counts were generated using featureCounts (v1.5.1). The mouse genome version mm10 was utilized.

### Publicly available datasets

Various publicly available datasets were used in this study to explore cell type-specific expression of differentially expressed genes. Published single cell data [89] were utilized to explore neuron-, astrocyte-, and microglia-specific expression of genes. Immunome-related genes were retrieved from the Immunome database. The Immune Response In Silico (IRIS) dataset was used to explore immunity-related genes [90] [91]

### Primary microglia cultures

Primary mouse microglia cell cultures were prepared as previously described for wild-type pups [92]. In brief, newborn mice (P1 pups) were used to prepare mixed glia cultures. Cells were grown in DMEM (Thermo Fisher Scientific) with 10% FBS, 20% L929 conditioned medium and 100 U ml–1 penicillin– streptomycin (Thermo Fisher Scientific). Microglia were collected 10-12 days after cultivation by shake off, counted and plated in DMEM supplemented with 10% FBS, 20% L929 conditioned medium and 100 U ml–1 penicillin–streptomycin. The microglia were shaken off up to two times.

### Ex-vivo isolation of microglia

PFC regions were dissected, mechanically dissociated and digested for 15 minutes with liberase (0.4 U/mL; Roche) and DNAse I (120 U/mL; Roche) at 37°C. Subsequently, the cell suspension was passed through a 70 µm cell strainer. Myelin debris was eliminated by the Percoll density gradient. Single cell suspension was labeled by using anti-mouse CD45 BV 421 (Clone 30-F11, Biolegend) and CD11b FITC (Clone M1/70, Biolegend). Antibody-labeled CD45^low^ CD11b^+^ microglial cells were sorted using a FACSAria 4L SORP cell sorter (Becton Dickinson) The purity of the sorted microglial cells was above 90%.

### Primary neuronal culture

Primary neuronal cultures were prepared from E17 pregnant mice of CD1 background (Janvier Labs, France). Briefly, mice were sacrificed and the brains of embryos were taken out, meninges removed, and the cortex dissected out. The cortexes were washed in 1× PBS (Pan Biotech, Germany). Single-cell suspensions were generated by incubating them with trypsin and DNase before careful disintegration. One hundred and thirty thousand cells per well were plated on poly-D-lysine-coated 24-well plates in Neurobasal medium (Thermo Fisher Scientific, Germany) supplemented with B-27 (Thermo Fisher Scientific, Germany). Primary cortical neurons were used for experiments at DIV10-12.

### Cell lines

All human iPSCs used in this study are commercially available and reported to be derived from material obtained under informed consent and appropriate ethical approvals.

### Differentiation of microglia from induced pluripotent stem cells

Human induced pluripotent stem cells lines (hiPSCs) (Cell line IDs: KOLF2.1J [93] were obtained from The Jackson Laboratory; BIONi010-C and BIONi037-A were both from the European bank for Induced Pluripotent Stem Cells) were differentiated to microglia as previously described [94]. In brief, 3 × 10^6 iPSCs were seeded into an Aggrewell 800 well (STEMCELL Technologies) to form embryoid bodies (EBs), in mTeSR1 and fed daily with medium plus 50 ng/ml BMP4 (Miltenyi Biotec), 50 ng/ml VEGF (Miltenyi Biotec), and 20 ng/ml SCF (R&D Systems). Four-day EBs were then differentiated in 6-well plates (15 EBs/well) in X-VIVO15 (Lonza) supplemented with 100 ng/ml M-CSF (Miltenyi Biotec), 25 ng/ml IL-3 (Miltenyi Biotec), 2 mM Glutamax (Invitrogen Life Technologies), and 0.055 mM beta-mercaptoethanol (Thermo Fisher Scientific), with fresh medium added weekly. Microglial precursors emerging in the supernatant after approximately 1 month were collected and isolated through a 40 um cell strainer and plated in N2B27 media supplemented with 100 ng/ml M-CSF, 25 ng/ml interleukin 34 (IL-34) for differentiation.

### Quantitative PCR experiment

cDNA synthesis was performed using the miScript II RT Kit (Qiagen, Germany) according to the manufacturer’s protocol. In brief, 200 ng total RNA was used for cDNA preparation. HiFlex Buffer was used so that the cDNA could be used for both mRNA and microRNA quantitative PCR (qPCR). A microRNA-specific forward primer and a universal reverse primer were used for quantification. The U6 small nuclear RNA gene was employed as an internal control. For mRNA quantification, gene-specific forward and reverse primers were used. The relative amounts of mRNA were normalized against GAPDH. The fold change for each microRNA and mRNA was calculated using the 2–ΔΔCt method^18^. The Light Cycler® 480 Real-Time PCR System (Roche, Germany) was used to perform qPCR.

### Caspase 1 activation assay

Caspase-Glo® 1 Inflammasome Assay (Promega, Germany) was used to detect caspase 1 activation as described in the manufacturer’s protocol. In brief, microglia, treated with ASO/inhibitor or primed with LPS and stimulated with ATP, were seeded on opaque, flat-bottom 96-well plates (Cellstar, Germany) at 50,000 per well in 100 ul DMEM supplemented with 10% FBS, 20% L929 conditioned medium and 100 U ml–1 penicillin–streptomycin. 100 ul of Caspase-Glo buffer was mixed with cell medium. Plates were incubated at room temperature for 1 h. Luminogenic caspase activity was measured using a FLUOstar Omega plate reader (BMG Labtech).

### Microglia phagocytosis assay

The microglia phagocytosis assay was performed as described^19^. Primary microglia cultures were plated at a density of 18 × 10^4^ in poly-D-lysine-coated 24-well plates in DMEM supplemented with 10% FBS, 20% L929 conditioned medium and 100 U ml–1 penicillin–streptomycin. Immortalized microglia (IMG) cultures were plated at a density 5 × 10^3^ in poly-D-lysine-coated 24-well plates in DMEM supplemented with 10% FBS, 1X Glutamine (Millipore), and 100 U ml–1 penicillin–streptomycin. To evaluate phagocytosis, treated microglia were incubated with fluorescent latex beads of 1 μm diameter (green, fluorescent 496/519; Sigma-Aldrich) for 1 h at 37°C, rinsed, and fixed with 4% formaldehyde. Cells were stained using the Iba1 (CD68) antibody (1:500; Wako) and DAPI. A confocal microscope was used for imaging at a low magnification (10x). ImageJ was used to quantify fluorescent latex beads. Region of interests (ROIs) were selected as microglial cells outlined with the Iba1 immunostaining to quantify beads. An intracellular section of the cell was selected to assure engulfment of latex beads by microglia. Similar acquisition parameters were used for each individual experiment. The results were expressed as the percentage of phagocytic index (# of total engulfed beads in an image / # of total cells identified in an image; n = 13 independent experiments).

### Synaptic pruning in primary microglia neural co-culture

Primary cortical neurons were seeded at a density of 130,000 on poly-D-lysine-coated 13 mm coverslips in 24-well plates in Neurobasal medium supplemented with B-27. Primary cortical neurons were used for experiments at DIV10-12. Treated primary microglia cultures were harvested from T-75 flasks and 4000 cells were seeded to each neural culture well. Plates were kept at 37°C for three days. On the third day, the cells were washed and fixed with 4% PFA (Sigma Aldrich, Germany) and 100 mM NH4Cl (Merck, Germany) respectively, at room temperature for 30 minutes. Next, the cells were washed in permeabilization and blocking buffer (0.1% Triton-X [Merck, Germany] + 3% bovine serum albumin (BSA) [AppliChem GmbH, Germany]) on a shaker. The cells were then incubated with primary antibodies for 1 hour at room temperature. The antibodies used included synaptophysin 1 (guinea pig, SySy), PSD-95 (rabbit, Cell Signaling,), and Iba1 (goat, Abcam). After incubation, the cells were washed in PBS and then incubated with a secondary antibody for 1 hour at room temperature. As secondary antibodies, Cy3 (donkey, anti-guinea pig, Jackson Imm.), Abberrior STAR 635p (goat, anti-rabbit) were used. Mowiol (Merck, Germany) and DAPI were used as a mounting medium. Images were taken with a multicolor confocal STED microscope (Abberior Instruments GmbH, Göttingen, Germany). Analysis of colocalization of pre- and post-synaptic markers were performed using SynQuant plugins in Fiji (v 2.0.0).

### Dendritic spine analysis

As described above, primary cortical neurons and primary microglia were co-cultured and fixed with 4%PFA. Dendritic spines were labeled as described [95]. In brief, the cells were aspirated and 2-3 crystals of Dil stain (Life Technologies-Molecular Probes) were added to each culture well and incubated on a shaker for 10 minutes at room temperature. Cells were washed with PBS until no crystals were visible and incubated overnight at room temperature. On the following day, the cells were washed and mounted with Mowiol. For high-magnification images, a multicolor confocal STED microscope with a 60× oil objective was used. Spine density and total spine length were measured by using ImageJ.

### Protein extraction of primary microglia

Primary microglia cell lysates were used to detect ZBP1 in RIPA fractions. Primary microglia were seeded in a 6-well plate at a density of 1 × 10^6^ in each well. Cells were collected in a RIPA buffer supplemented with 1 × protease inhibitor. Samples were kept on ice for 15 minutes and vortexed every 5 minutes and then centrifuged at 5000 rpm for 15 minutes at 4°C before supernatants were transferred to a new tube and stored at –20°C. The protein concentration was measured using a BCA assay.

### Immunoblot analysis

For standard immunoblot analysis, 20 ug of samples were mixed with 1× Laemmli buffer (Sigma, Germany), heated for 5 min at 95°C and loaded onto 4–15% Mini-PROTEAN® TGX™ Precast Protein Gels (Bio-Rad, Germany). Proteins were transferred on nitrocellulose membranes and membranes were blocked with 5% BSA in PBS-Tween. Membranes were incubated with primary antibodies in 5% BSA in PBS-Tween. Fluorescent-tagged secondary antibodies (LI-COR) were used for visualization of proteins. Imaging was performed using a LI-COR ODYSSEY. HSP-70, GAPDH were used as a loading and run on the same gel.

### Treatment of microglia

Microglia activation by LPS was used as a positive control. For this, microglia cells were first primed with 100 ng/ml ultrapure LPS (E. coli 0111:B4, Invivogen) and then incubated at 37°C. After this, 5 mM ATP were added to the culture and incubated for 30 minutes. Caspase 1 assay and phagocytosis assay were performed from these cultures. For immunoblot, cell lysate was prepared. For miR99b-5p-related analysis, microglia were either treated for two days with miR99b-5p inhibitor/negative control or ASOs in T-75 after first harvesting or after harvesting cells were seeded in a 24-well culture plate.

### Luciferase assay

Seed sequences of miR-99b-5p and pairing 3ʹUTR sequences of Zbp1 were generated with TargetScan. Cloned 3ʹUTR sequence of Zbp1 and scrambles UTR were purchased from Gene Copoeia (https://www.genecopoeia.com/product/mirna-target-clones/mirna-targets/). UTR was cloned downstream to firefly luciferase of pEZX-MT06 Dual-Luciferase miTarget™ vector. The pEZX-MT06-scrambled UTR or pEZX-MT06-Zbp1 3ʹUTR construct and miR99b-5p mimic or negative control were co-transfected into HEK293-T cells cultured in 24-well plates using EndoFectin™ Max Transfection Reagents (Gene Copoeia) according to the manufacturer’s protocol. 48 hours after transfection, Firefly and Renilla luciferase activities were measured using a Luc-Pair™ Duo-Luciferase HS Assay Kit (for high sensitivity) (GeneCopoeia). Firefly luciferase activity and Renilla luciferase activity were normalized. The mean of luciferase activity and of Firefly/Renilla was considered for the analysis.

### Statistical analysis

Unless otherwise noted, statistical analysis was carried out with GraphPad Prism software version 8.0. Statistical measurement is shown as mean ± SD. Each n represents a biological sample. Either a two-tailed unpaired t-test or a two-way ANOVA with Tukey’s post hoc test were applied to analyze the data. Enriched gene ontology and pathway analysis was performed using Fisher’s exact test followed by a Benjamini-Hochberg correction.

## Conflict of interest

The authors declare no conflict of interest.

## Acknowledgment

This work was supported by the following grants to AF: The DFG (*Deutsche Forschungsgemeinschaft*) priority program 1738, SFB1286, the EPIFUS project, Germany’s Excellence Strategy - EXC 2067/1 390729940. FS was supported by the GoBIO project miRassay. Urs Heilbronner is supported by the European Union’s Horizon 2020 Research and Innovation Programme (PSY-PGx, grant agreement No 945151) and the Deutsche Forschungsgemeinschaft (DFG, German Research Foundation, project number 514201724). LE is supported by the Studienstiftung des Deutschen Volkes and the International Max-Planck Research School for The Mechanisms of Mental Function and Dysfunction (IMPRS-MMFD). DKV is supported by grants from the Chan Zuckerberg Initiative Neurodegenerative’s Challenge Network (2020-221779(5022) & 2021-235147). TS was supported by grants from the *Deutsche Forschungsgemeinschaft* (DFG; SCHU 1603/4-1, 5-1, 7-1), the German Ministry of Education and Research (BMBF; 01EE1404H), and the Dr. Lisa Oehler Foundation (Kassel, Germany). PF was supported by a grant from the DFG (DFG; FA 241/16-1)

## Author contributions

L.K., F.S., and A.F. designed research; L.K., M.R.I., J.Z., J.S., R.P., S.B., L.E., A.B., D.K.V., I.D., F.O., and A.F. performed research; U.H., A.B., M.B., F.S., M.O.K., E.C.S., M.S., E.Z.R., G.J., T.G.S., P.F. recruited and phenotyped patients; L.K., M.R.I., D.M.K., A.M., T.P., and A.F. analyzed data; and L.K., F.S., and A.F. wrote the paper with input from other co-authors.

## Notes

### Competing Interest Statement

The authors have declared no competing interest.

### Summary of Updates

Author contributions added.

