## Supplementary figures for "A novel miR-99b-5p-*Zbp1* pathway in microglia contributes to the pathogenesis of schizophrenia"

### Supplemental Figures 1-8

**Figure S1**

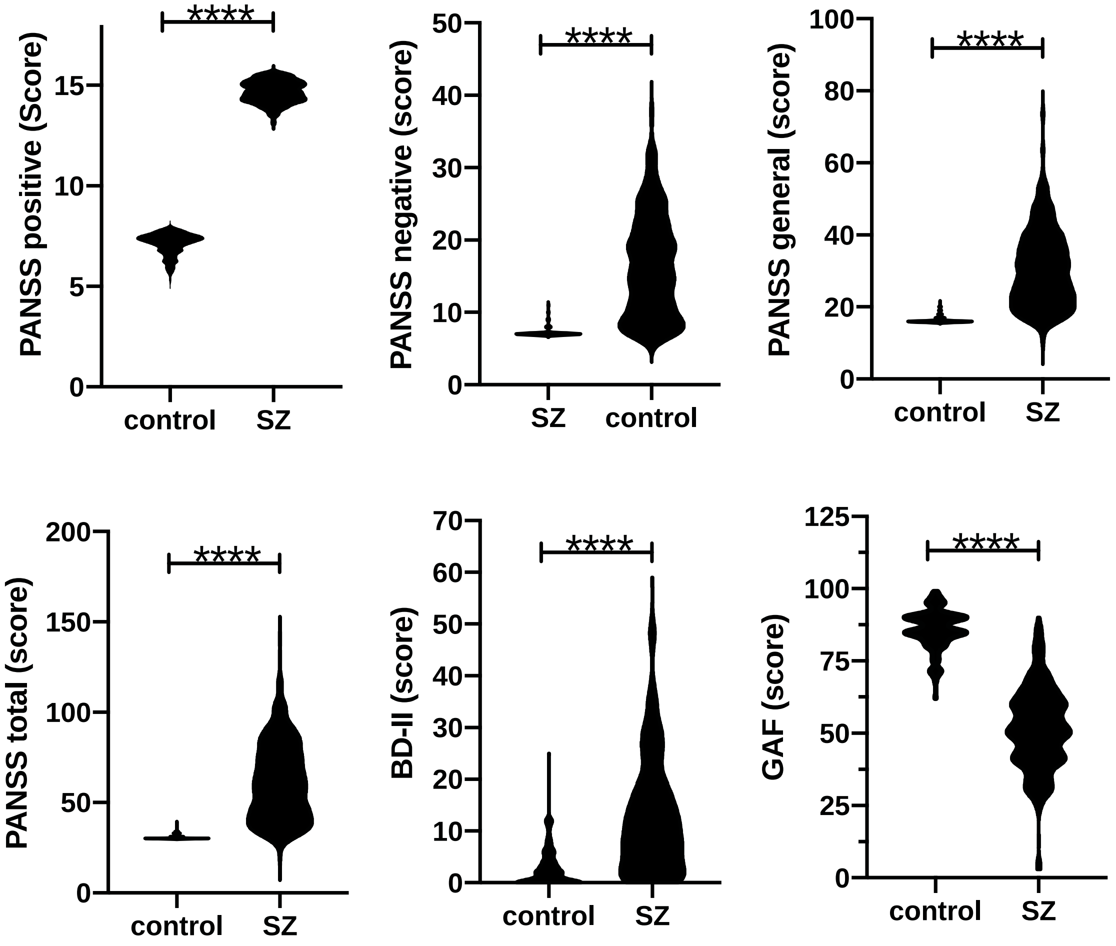


**Figure S1. Clinical phenotypes of the individuals subjected for smallRNA-seq analysis**. We analyzed 242 healthy controls and 331 SZ patients of the PsyCourse study. Depicted are the clinical phenotypes that differ significantly between groups, namely the positive and negative syndrome rating scale (PANSS), the total PANSS, the Beck depression inventory (BDI-II) and the global assessment of functioning (GAF) scores. ****P < 0,0001, *t*Test.

#### Figure S2

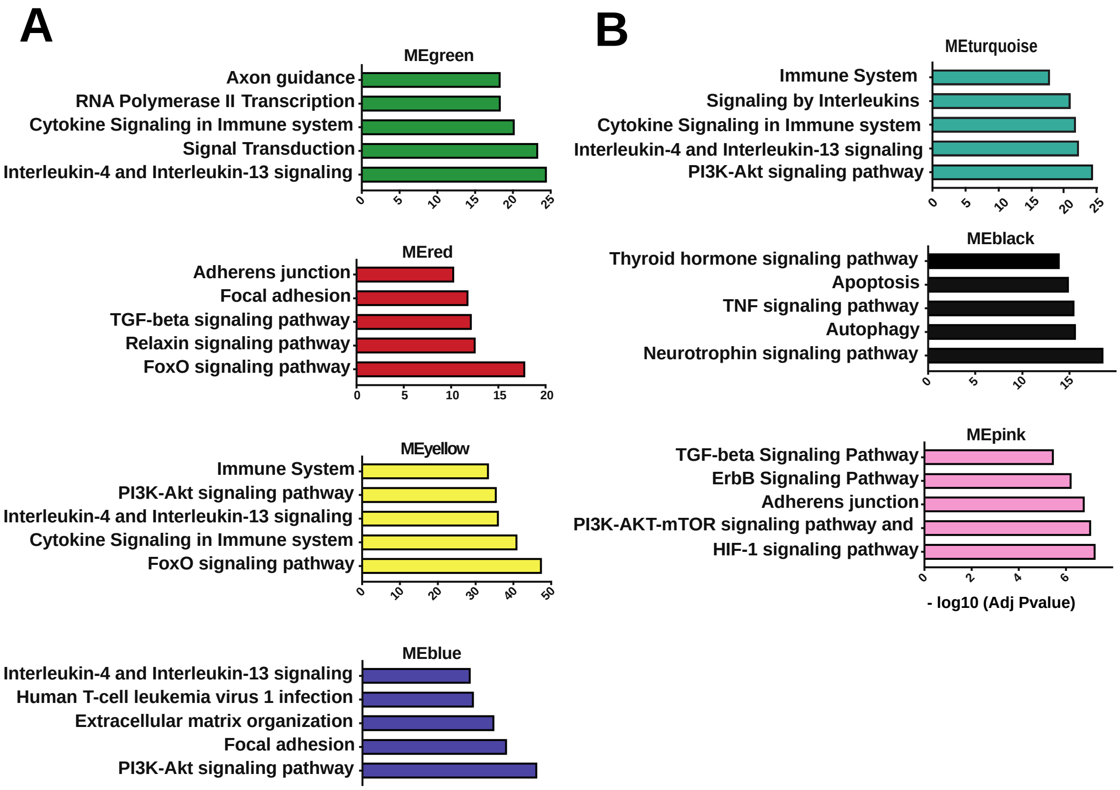


**Figure S2. GO-term analysis for confirmed mRNA targets of microRNAs found within the different co-expression modules**. Confirmed mRNA targets were identified for the microRNAs within each of the detected co-expression modules (see Fig 1b). The corresponding gene-lists were subjected to GO-term analysis. **A.** Bar graphs depicting the top 5 GO-terms for the 4 co-expression modules (MEgreen, MEred, MEyellow and MEblue) that were increased in schizophrenia patients. **B. A.** Bar graphs depicting the top 5 GO-terms for the 3 co-expression modules (MEturquoise, MEblack, and MEpink) that were decreased in schizophrenia patients.

**Figure S3**

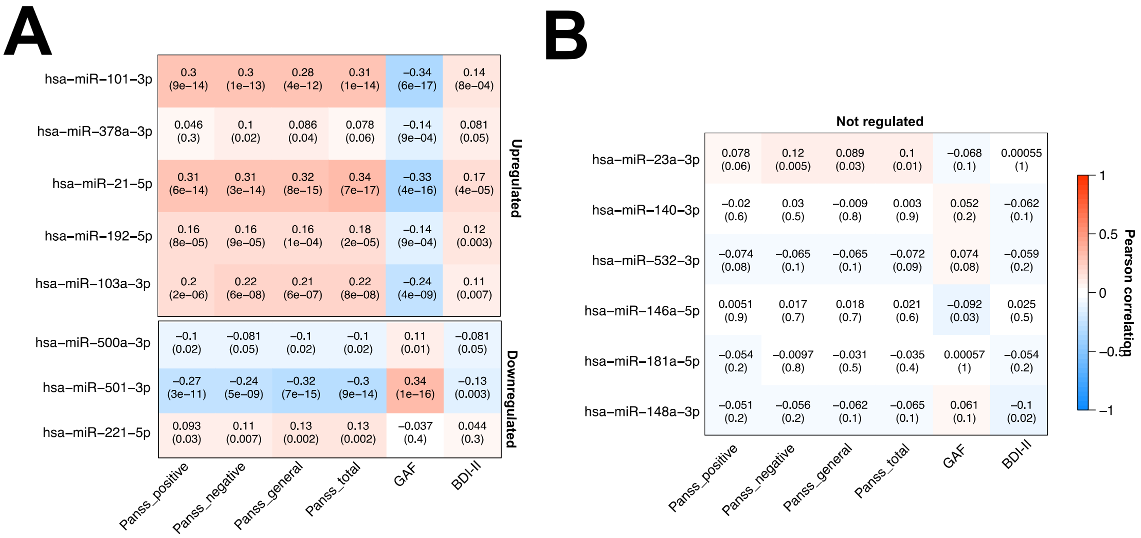


**Fig. S3. Correlation of candidate miR expression to SZ phenotypes. A.** Heat map showing the correlation of candidate miR expression levels of individuals of the PsyCourse study to the clinical phenotypes. The numbers in each rectangle represent the correlation (upper number) and the corresponding p-value (lower number). Values for miR-99b-5p are shown within Fig 1h. MiR-21-5p and miR-501-3p have been previously linked to SZ (Refs 21, 29) and show the highest correlation values. MiR-221-5p is significantly correlated to the PANSS scores but not the GAF and BDI-II. **B.** Heat map showing the correlation of six miRs from our dataset that were not differentially expressed in the blood or brain of SZ patients (not regulated). miR-23a-3p, miR-140-3p and miR-532-3p were randomly selected, while miR-146a-5p, miR-181a-5p and miR-148a-3p were recently identified as a biomarker signature for Alzheimer's disease (Ref 92).

**Figure S4**

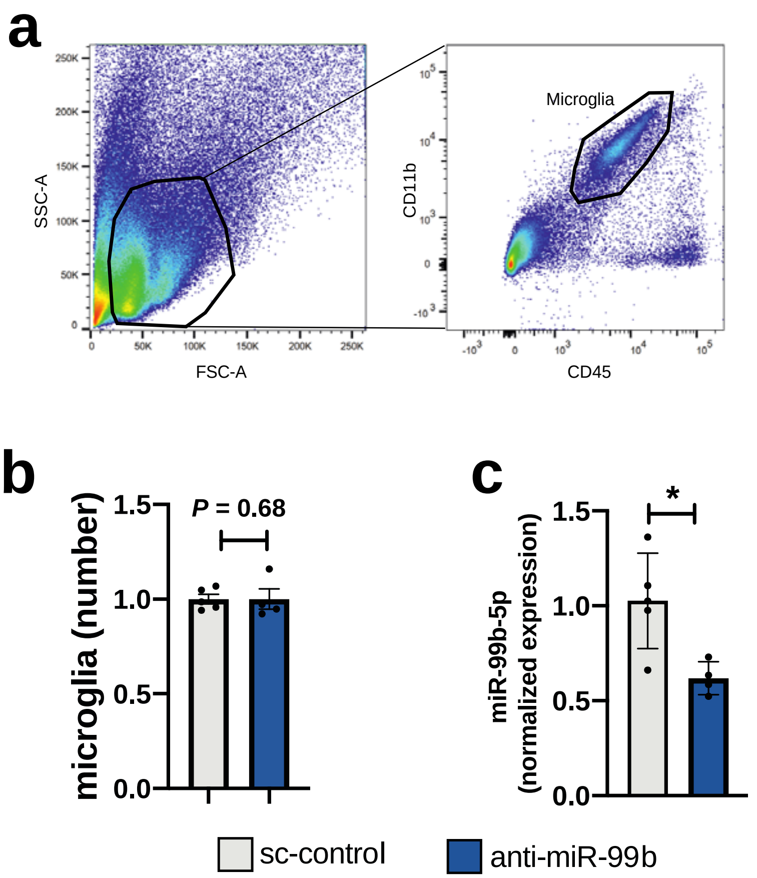


**Figure S4. Analysis of microglia obtained via FACS from the PFC of mice injected with anti-miR-99b or sc-control LNAs**. **a.** Dot plots depicting the gating strategy for the isolation of $CD45^{low}/CD11b^{+}$ microglial cells **b.** Bar graph showing the number of microglia (normalized values) obtained from the PCR of mice injected with either sc-control LNAs or anti-miR-99b. No statistical difference was observed between groups. **c.** Bar graph showing the expression of miR-99b-5p as determined via qPCR in PFC tissue isolated from mice injected into the PFC with sc-control LNAs or anti-miR-99b. Microglial miR-99b-5p levels are significantly impaired in mice that received anti-mIR-99b injection into the PFC. *$P$ < 0,05. Error bars indicate SEM.

**Figure S5**

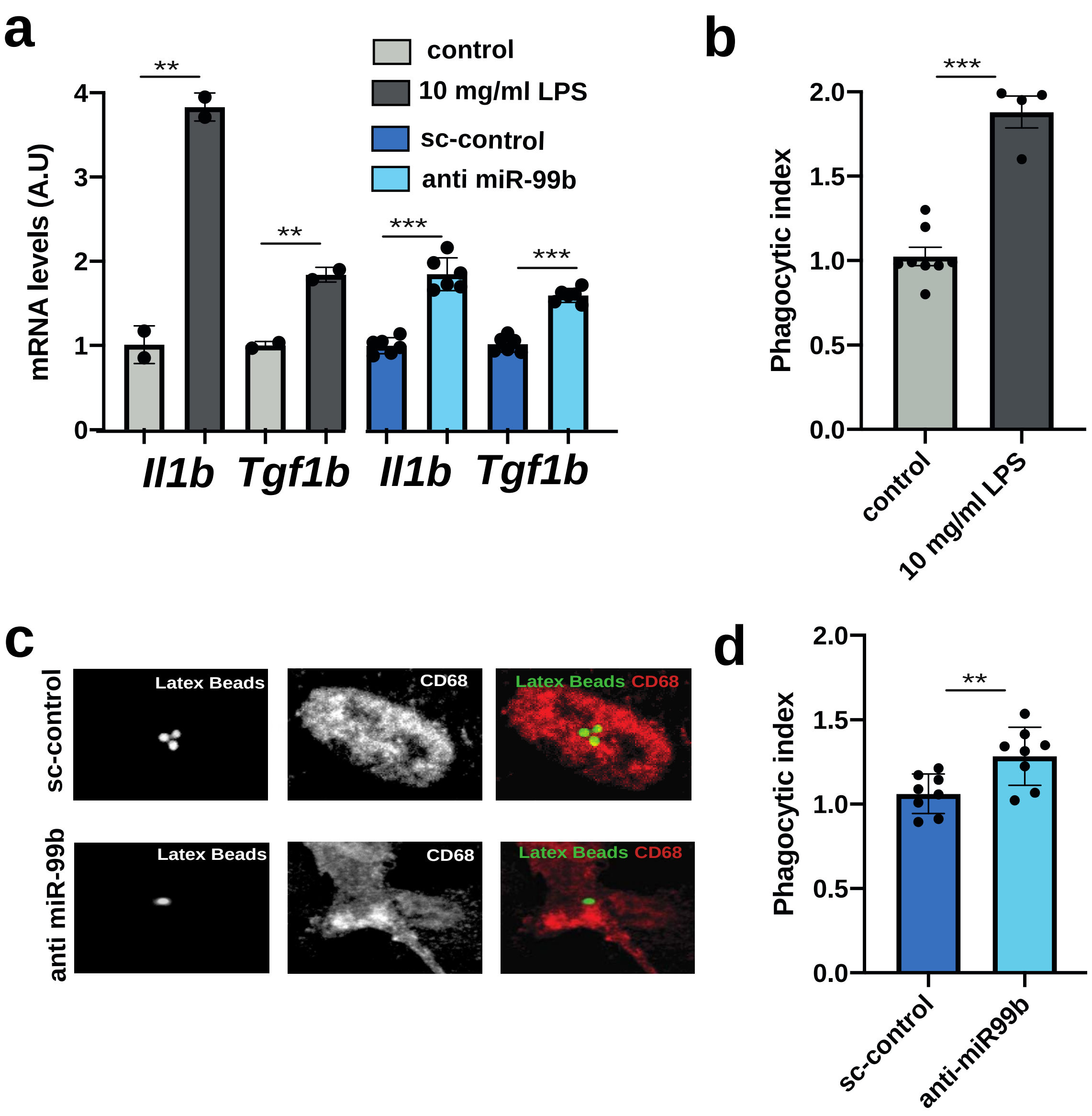


**Figure S5. Loss of miR-99b-5p levels increases the expression of pro-inflammatory cytokines and phagocytosis in IMG cells**. **a**. IMG cells were treated with vehicle control solution (control) or 10mg/ml LPS. In a similar experiment, IMG cells were treated with LNPs loaded with either sc-control LNAs or anti-miR-99b. qPCR analysis was performed for the pro-inflammatory cytokines *Il1b* and *Tgf1b*. The expression of *Il1b* and *Tgf1b* were significantly increased upon LPS treatment or anti-miR-99b treatment when compared to the respective control groups. **b.** Bar graph showing that the phagocytic index increases in IMG cells upon LPS treatment. **c.** Representative images showing the uptake of latex beads by IMG cells treated with either sc-control LNAs or anti-miR-99b. **d.** Bar graph showing the quantification of (c). Treatment with anti-miR-99 increases the phagocytic index. **P* < 0,05, ***P* < 0,01, ****P* < 0,001. Error bars indicate SEM.

**Figure S6**

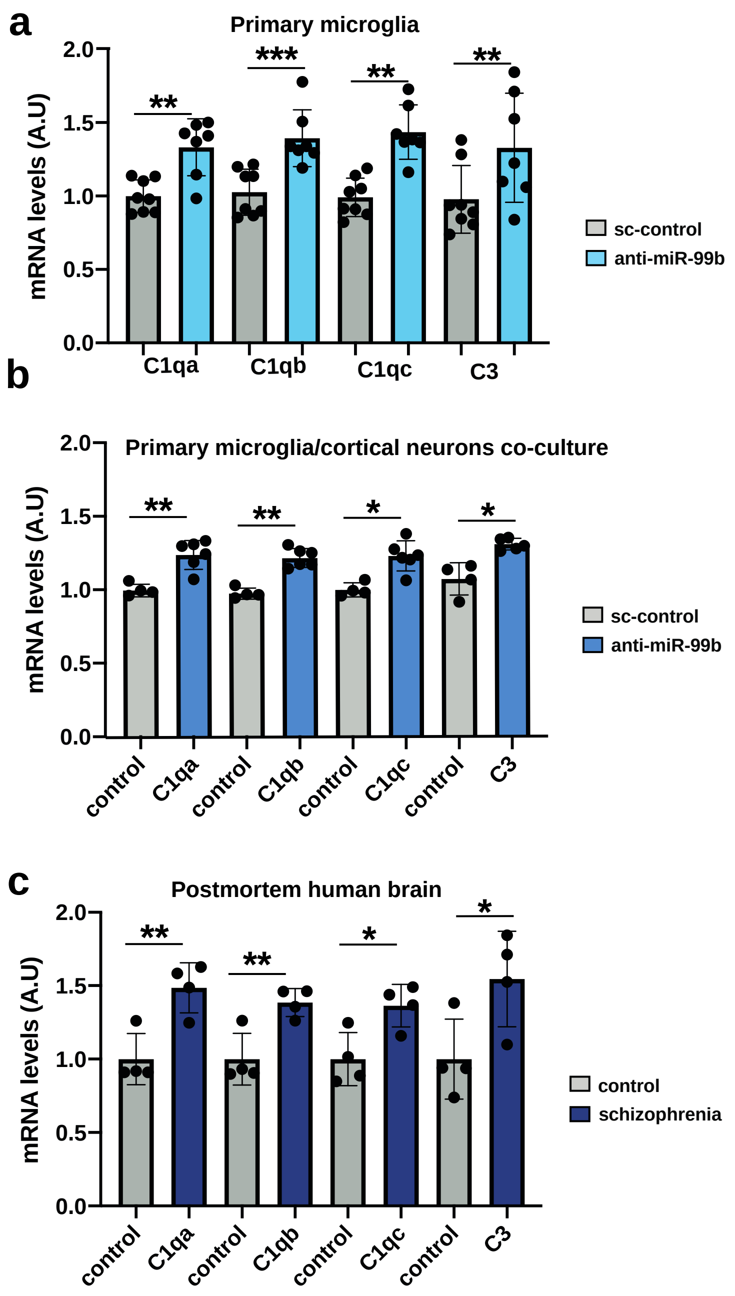


**Figure S6. Genes linked to innate immunity and synaptic pruning are increased in microglia treated with anti-miR-99b and in schizophrenia patients. a.** Bar graph showing qPCR results for *C1qa*, *C1qb*, *C1qc*, and *C3* in primary microglia treated with sc-control LNAs or anti-miR-99b. **b.** Bar graph showing qPCR results for *C1qa*, *C1qb*, *C1qc*, and *C3* when RNA was isolated from co-cultures in which primary cortical neurons were treated with microglia that had received LNPs loaded with either sc-control LNAs or anti-miR-99b. c. QPCR analysis was used to measure *C1qa*, *C1qb*, *C1qc*, and *C3* expression in human postmortem PFC samples obtained from control individuals and SZ patients (*t*Test, $^{**}P < 0{,}01$, $^{*}P < 0{,}05$; n = 4/group). Error bars indicate SEM.

**Figure S7**

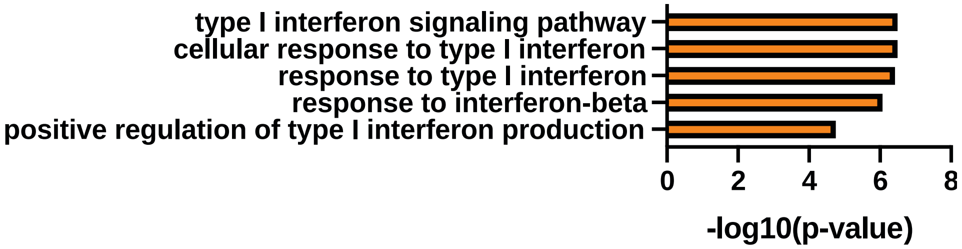


**Figure S7. GO-term analysis of miR-99b-5p target genes up-regulated in the PFC of mice treated with anti-miR99b.** Bar graph depicting the 5 TOP GO-terms when analyzing miR-99b-5p target genes that were up-regulated in the PFC cortex of mice injected to the PFC with anti-miR-99b.

**Figure S8**

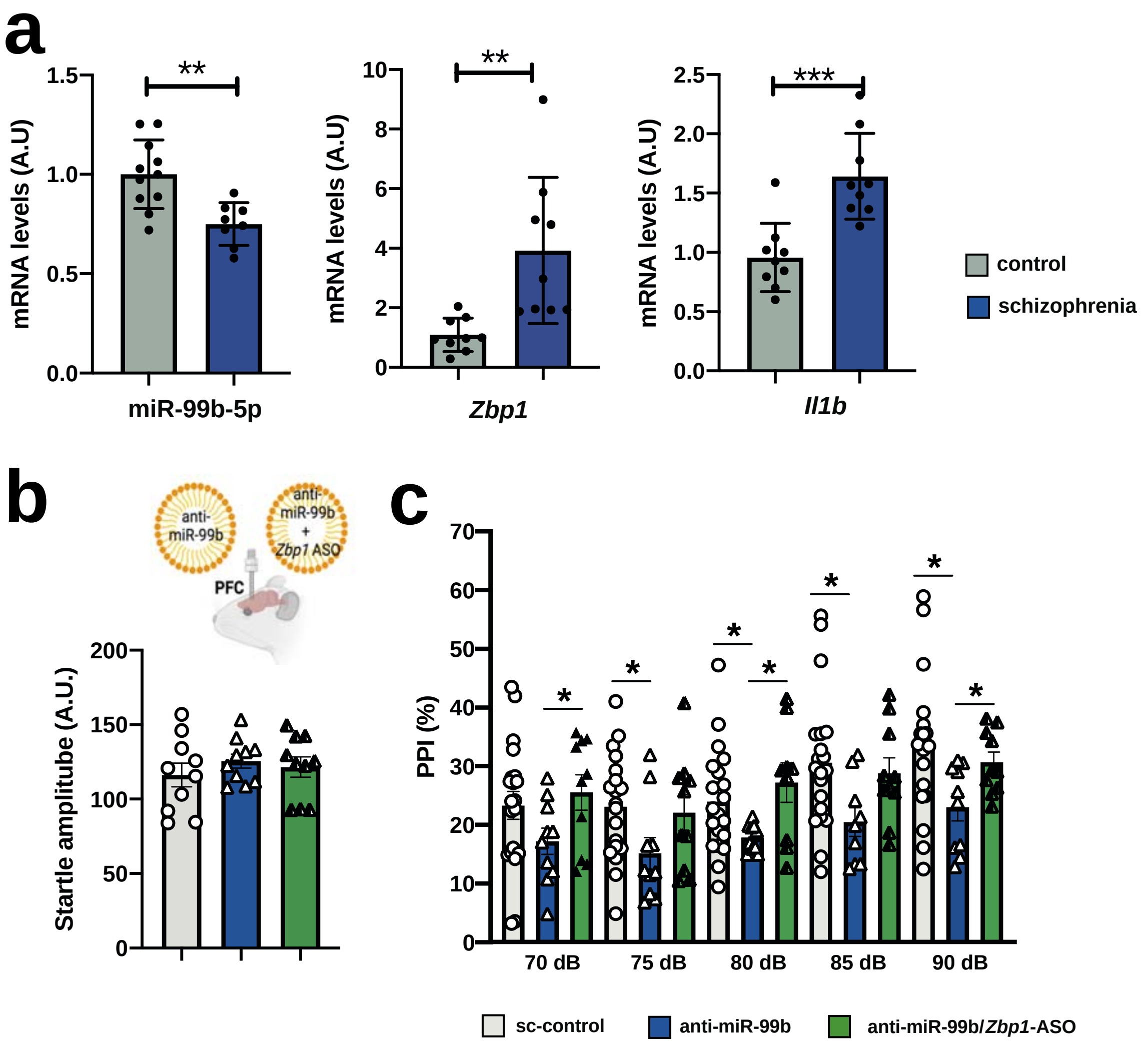


**Figure S8. Anti-miR-99b mediated impairment of PPI depends on *Zbp1*. a.** Bar graph showing qPCR results for miR-99b-5p, *Zbp1* and *Il1b* in PFC tissue of SZ patients and control individuals. While miR-99b-5p levels are decreased, expression of the miR-99b-5p target gene *Zbp1* is increased in SZ patients. In line with this expression of the pro-inflammatory cytokine *Il1b*, which is regulated by *Zbp1*, is increased in SZ patients when compared to control. (*t*Test, ****P* < 0,001; ***P* < 0,01, **P* < 0,05; n = 9/group) **b.** Upper panel: Experimental design. LNPs loaded with either sc-control, anti-miR-99b or anti-miR-99b together with *Zbp1* ASOs were injected into the PFC before mice were subjected to behavioral testing. Lower panel: Bar graph showing that the basic startle response is not different amongst experimental groups (n = 10 / group). **c.** Bar graph showing results from the PPI experiment comparing mice injected to the PFC with either sc-control (n = 10), anti-miR-99b (n = 10) or anti-miR-99b together with *Zbp1* ASOs (n = 10). One-way ANOVA revealed a significant difference amongst groups (P < 0,0001, $F_{(1, 19)}$ = 27). In agreement with our previous data, PPI is impaired when comparing the sc-control group to mice injected with anti-miR-99b (P < 0,0001, $F_{(4, 89)}$ = 27; Two-Way ANOVA). In contrast, no difference was observed when the sc-control group was compared to mice injected with anti-miR-99b together with *Zbp1* ASOs (P = 0,3946, $F_{(4, 89)}$ = 0,7).*t*Test was used to compare groups at the indicated dB values. Asterisks indicate significance (*$P <$ 0,05). Error bars indicate SEM.
